# FULL-seq reveals isoform-specific regulatory architecture in single cells

**DOI:** 10.64898/2026.11.17.767299

**Authors:** Mingchao Wang, Jianhuan Qi, Francesco Monticolo, Xingjian Chen, Amanda Qiao, Lee Zou, Jian Shu

## Abstract

Gene regulation is commonly modeled at the gene level, yet genes express multiple isoforms from distinct transcription start sites whose regulatory architecture is not resolved by current approaches. Here we introduce FULL-seq, a scalable single-cell multiomic platform that jointly profiles chromatin accessibility and full-length transcript coverage from the same cell, enabling direct coupling of regulatory elements to alternative first exon (AFE) usage. FULL-seq achieves high sensitivity for both transcriptome and chromatin accessibility while providing transcript-wide coverage across mRNAs up to 15 kb with precise exon-level mapping. We uncover diverse promoter–AFE regulatory architectures, revealing distinct transcription factor programs across alternative first exons of the same gene, and show that gene-level annotation frameworks can obscure regulatory relationships in multi-promoter genes. Applying FULL-seq to human cancer cells further reveals metastasis-associated AFE programs that are not apparent from gene-level expression. These AFE programs were reproduced in independent colorectal cancer cell-line datasets, and CALB1 AFE activation was observed in a subset of liver metastases from an independent patient cohort. Furthermore, CRISPRi-mediated repression of metastasis-associated CALB1 AFEs reduced colorectal cancer cell migration. Together, our findings establish alternative promoter and first-exon usage as a functionally relevant layer of gene regulation and demonstrate that FULL-seq enables regulatory relationships and cellular phenotypes to be connected at the promoter–isoform level.

## Main

Gene regulation is commonly conceptualized at the gene level, where chromatin accessibility at promoters is assumed to directly govern transcriptional output^1–5^. However, most mammalian genes generate multiple transcript isoforms through alternative transcription start sites (TSSs), resulting in distinct alternative first exons (AFEs)^6,7^. Unlike alternative splicing events generated after transcription initiation, AFE usage is directly coupled to promoter selection and therefore provides a particularly direct interface between chromatin regulation and transcript isoform identity. These isoforms are often clinically relevant^7^, tissue-specific^6^, and dynamically regulated^8^. This raises a fundamental question: how chromatin accessibility controls transcription at the level of individual isoforms rather than aggregated gene output?

Despite recent advances in single-cell multiomic technologies that jointly profile chromatin accessibility and gene expression^1,9–12^, current approaches remain fundamentally limited to gene-level resolution. Transcript reads from different isoforms are typically aggregated into a single expression measurement, while accessible chromatin regions are assigned to genes using canonical promoter annotations or genomic proximity^13,14^. For genes with multiple transcription start sites, such aggregation can obscure promoter-specific regulation and incorrectly assign regulatory elements to transcript outputs. Consequently, distinct isoforms originating from the same gene may be controlled by different regulatory programs while appearing indistinguishable in conventional gene-level analyses.

Methods that improve transcript-structure resolution provide important complementary information. Full-length single-cell RNA-sequencing approaches^15–17^ can resolve transcript structures but generally lack matched chromatin accessibility from the same cell. More recently, approaches such as ScISOr-ATAC^18^ have enabled joint profiling of chromatin accessibility and transcript isoform structure, providing new opportunities to investigate exon-level regulation. However, targeted transcript coverage, limited per-cell molecular depth, and incomplete transcription start site (TSS) annotation constrain genome-wide quantitative analysis of promoter-resolved isoform regulation. Thus, directly connecting accessible regulatory elements with full-length transcript outputs at single-cell resolution remains challenging.

Here, we introduce **FULL-seq** (**full**-length single-cell RNA and ATAC **seq**uencing), a scalable short-read single-cell multiomic platform that jointly profiles chromatin accessibility and transcript-wide RNA coverage from the same cell. By linking accessible chromatin and full-length cDNA through shared cellular barcodes, FULL-seq directly resolves regulatory relationships between accessible promoters and alternative first exon usage. We identify diverse promoter–AFE regulatory architectures and distinct transcription factor programs across alternative first exons of the same gene, revealing regulatory relationships that are obscured by gene-level analyses. Extending FULL-seq to human cancer models reveals cancer-state-specific AFE programs, including metastasis-associated AFEs that are undetectable at the gene-expression level and reproducible in independent colorectal cancer datasets. Finally, CRISPRi-mediated perturbation of metastasis-associated *CALB1* AFEs alters colorectal cancer cell migration, demonstrating that isoform-resolved regulatory differences identified by FULL-seq can have measurable phenotypic consequences. Together, these findings establish promoter–isoform regulation as an important dimension of transcriptional control and provide a framework for connecting chromatin state, transcript identity, and cellular phenotype at isoform resolution.

### Development of FULL-seq for isoform-resolved chromatin–transcriptome profiling

Alternative transcription start site usage generates transcript isoforms with distinct first exons and promoter contexts, representing an important mechanism for regulating gene expression across cell states. Because each alternative first exon is typically associated with a distinct promoter, understanding the regulatory logic of alternative transcription requires linking chromatin accessibility to isoform-specific transcription start sites.

Current single-cell multiomic approaches jointly profile chromatin accessibility and gene expression but typically operate at the gene level, aggregating RNA reads from all transcript isoforms into a single unit and linking chromatin accessibility to annotated transcription start sites. As a result, these methods cannot resolve regulatory relationships associated with alternative transcription start sites and isoform-specific promoters. Consequently, regulatory elements controlling alternative transcript isoforms may be obscured or incorrectly assigned when analyses are performed at gene resolution, introducing systematic bias (Extended Data Fig. 1a).

To overcome these challenges, we developed FULL-seq, which jointly profiles chromatin accessibility and high-quality full-length transcript coverage, achieving predominantly exonic mapping with minimal intronic noise. The workflow involves reverse transcription, Tn5 tagmentation, and single-cell barcoding, followed by physical separation of RNA and DNA fractions for independent library construction and sequencing (Fig. 1a). In this process, mRNA is reverse-transcribed into full-length cDNA before simultaneous Tn5-based tagging of accessible chromatin, allowing both transcriptome and chromatin accessibility profiles to be captured from the same cell. A key step in FULL-seq is reverse transcription using biotinylated dNTPs. Among tested variants, biotin-dCTP gave strong cDNA yield without impairing efficiency (Supplementary Fig. 1). Together, these technical and operational advantages position FULL-seq as a broadly accessible, high-resolution platform for full-length single-cell transcriptome and epigenome profiling.

**Fig.1:**
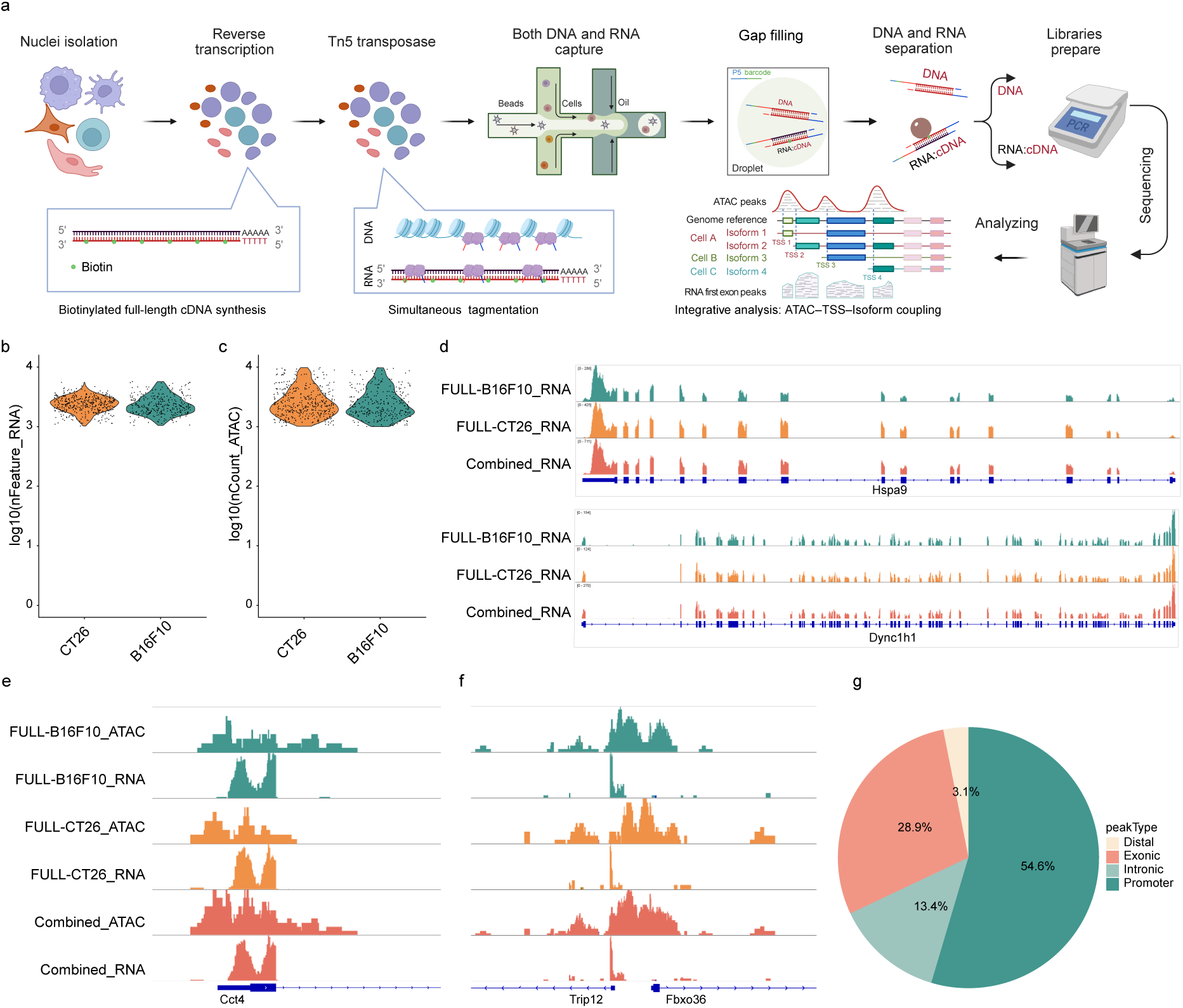
Overview and performance benchmarking of FULL-seq. **a,** Schematic of the FULL-seq workflow, highlighting nuclei isolation, biotinylated full-length cDNA synthesis, Tn5 tagmentation, droplet-based barcoding, RNA–DNA separation, and joint RNA/ATAC library preparation. **b,** Distribution of detected genes across CT26 and B16F10 cell lines. **c,** Distribution of ATAC read counts across CT26 and B16F10 cell lines. **d,** IGV browser tracks showing aggregated RNA coverage from CT26, B16F10, and their combined dataset at representative loci spanning 2.5 kb (top) and 15 kb (bottom). **e,** Integrated IGV browser tracks showing matched RNA and ATAC coverage from CT26, B16F10, and their combined dataset at representative loci. **f,** Integrated IGV browser tracks showing matched RNA and ATAC coverage at representative loci with multiple promoter–AFE pairs. **g,** Pie chart showing the distribution of conventional genomic annotations of ATAC peaks overlapping AFE regions. Peaks were classified as promoter, exonic, intronic, or distal using ArchR’s gene-based annotation framework.

To benchmark the performance of FULL-seq, we applied the method to CT26 (murine colorectal carcinoma) and B16F10 (murine melanoma) cells (n = 751 high-quality cells) and first evaluated RNA data quality. FULL-seq detected an average of approximately 2,500 genes per cell (Fig. 1b), demonstrating sensitivity comparable to conventional scRNA-seq. The resulting transcriptomic profiles clearly separated CT26 and B16F10 cells into distinct clusters (Extended Data Fig. 1b–d), confirming the robustness of gene expression measurements.

We next examined the quality of chromatin accessibility profiles generated by FULL-seq. The ATAC modality displayed clear nucleosome periodicity (Extended Data Fig. 2a) and strong enrichment of accessibility signals around transcription start sites (Extended Data Fig. 2b), two hallmarks of high-quality ATAC-seq data. Consistent fragment count distributions across CT26 and B16F10 cells (Fig. 1c), together with accurate separation of the two cell types based on ATAC profiles (Extended Data Fig. 2c–e), further demonstrate the reliability of chromatin accessibility measurements.

Furthermore, FULL-seq enables robust detection of full-length transcript coverage up to 15,000 bp while maintaining precise exon-level mapping across the entire transcript, consistently across cell populations (Fig. 1d).

Aggregated genome browser views further revealed striking co-localization between 5′ RNA transcription peaks and nearby ATAC peaks (Fig. 1e), clearly resolving multiple promoter–AFE pairs within individual loci (Fig. 1f). This spatial proximity between transcription start sites and accessible chromatin highlights the regulatory coupling between promoters and alternative first exons.

Finally, we examined the genomic annotations of chromatin accessibility peaks associated with alternative first exons (AFEs). Conventional peak annotation approaches assigned only a subset of these peaks to promoter regions, whereas a substantial fraction were labeled as intronic, exonic, or distal elements (Fig. 1g). This observation highlights the ambiguity inherent in gene-level peak annotation strategies and suggests that regulatory elements controlling alternative first exon usage may frequently reside outside canonical promoter regions.

Together, these results demonstrate that FULL-seq enables robust simultaneous profiling of full-length transcript isoforms and chromatin accessibility at single-cell resolution, providing a foundation for systematic analysis of the regulatory architecture underlying alternative transcription start site usage.

### FULL-seq improves transcript structure resolution and enables identification of alternative first exons

To evaluate the transcriptome profiling performance of FULL-seq relative to existing single-cell platforms, we compared RNA coverage patterns with those obtained from 10x snRNA-seq, 10x scRNA-seq, and 10x Multiome-seq. Aggregated genome browser tracks at representative loci revealed that FULL-seq produced continuous full-length RNA coverage across gene bodies, whereas 10x-based methods showed strong enrichment near transcript ends (Fig. 2a). More interestingly, FULL-seq maintained clean exon mapping even at log scale, where 10x data displayed prominent intronic and intergenic reads (Fig. 2a and Extended Data Fig. 3a-c). This difference was evident across genomic regions spanning both short (0.5 kb) and long (15 kb) loci. Consistent with these observations, >85% of FULL-seq reads mapped to exonic regions compared to 10x (Fig. 2b), despite using nuclei as input material, which often increases intronic retention in conventional methods. In addition, comparison with VASA-seq further highlighted FULL-seq’s advantage: VASA-seq exhibited pronounced intronic signal noise (Extended Data Fig. 3d), consistent with their previous reports^17^.

**Fig.2:**
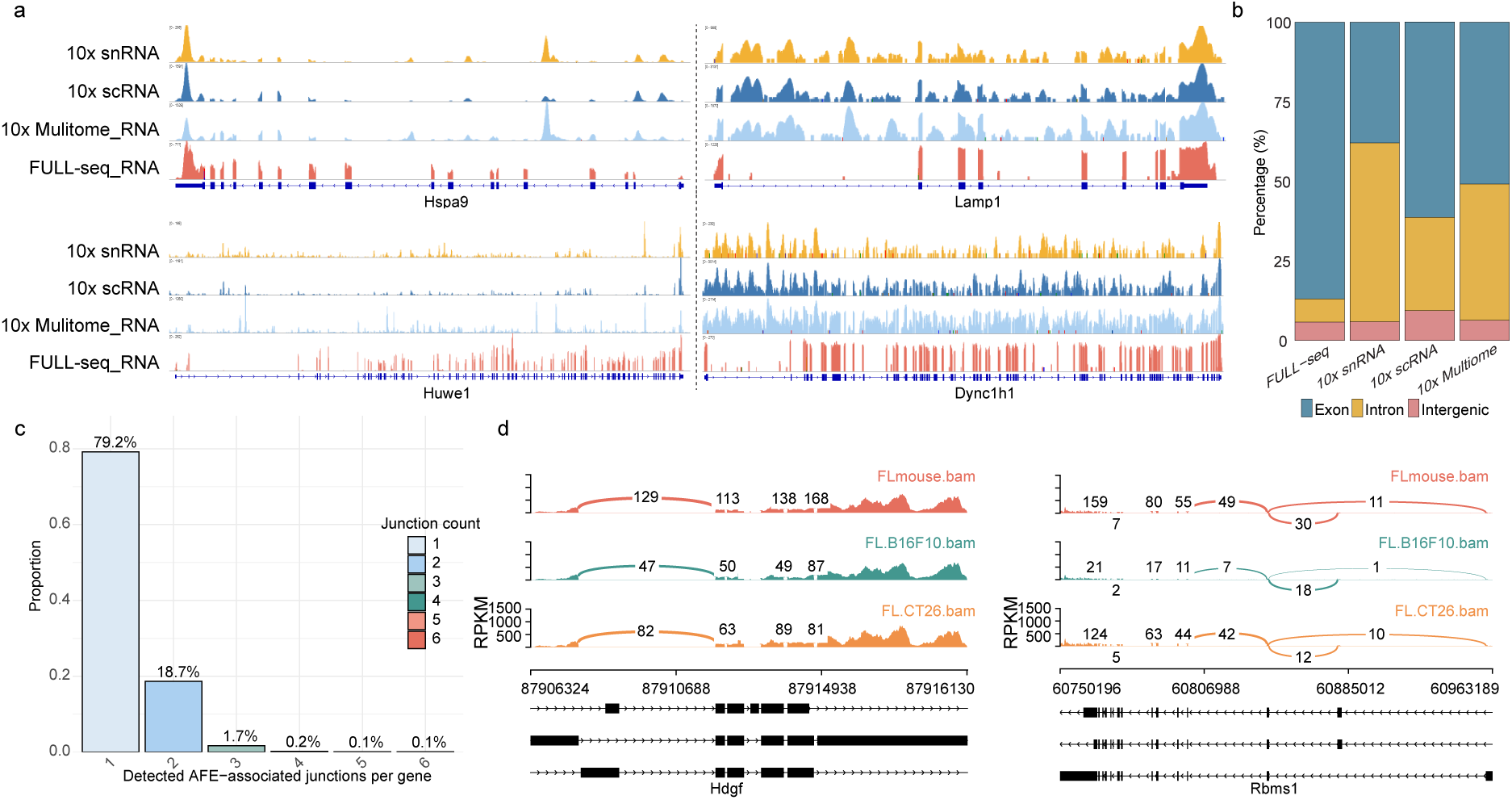
FULL-seq improves transcript structure resolution and enables identification of alternative first exons. **a,** IGV browser tracks comparing aggregated RNA coverage from FULL-seq and conventional platforms (10x snRNA-seq, 10x scRNA-seq, and 10x Multiome-seq) at representative loci, spanning 2.5 kb (top) and 15 kb (bottom) regions (linear scale, left; log scale, right). **b,** Fraction of reads mapped to exonic, intronic, and intergenic regions for each platform, showing the high exonic capture efficiency of FULL-seq. **c,** Distribution of the number of distinct detected AFE-associated junctions per gene in the pooled CT26 and B16F10 dataset. Among genes with at least one detected AFE-associated junction, 79.2% exhibited a single junction and 20.8% exhibited multiple junctions. **d,** Representative Sashimi plots of a mono-AFE gene (left) and a multi-AFE gene (two AFEs; right) across the combined sample as well as separately analyzed CT26 and B16F10 cells. Read density is shown as bar heights, and splice junction counts are indicated by connecting arcs.

The enhanced transcript coverage enabled annotation-guided analysis of alternative first exons (AFEs) across the transcriptome. In the pooled CT26 and B16F10 dataset, 79.2% of genes with detectable AFE-associated junctions exhibited a single distinct junction, whereas 20.8% exhibited multiple junctions (Fig. 2c). Similar distributions were observed when CT26 and B16F10 cells were analyzed separately (Extended Data Fig. 4a). Representative IGV analysis further revealed six distinct annotated AFEs within an individual gene, supported by RNA coverage and splice junction evidence (Extended Data Fig. 4b), illustrating the ability of FULL-seq to capture diverse first-exon-associated transcriptional patterns.

To better explore the regulatory logic of AFE usage, we categorized genes with multiple annotated AFEs into two groups based on isoform expression: (1) mono-AFE genes, which express a single AFE isoform, and (2) multi-AFE genes, which express two or more AFE isoforms. Sashimi plot visualization of representative mono-AFE genes confirmed specific AFE with clean exon–intron boundaries (Fig. 2d, left). We further examined multi-AFE genes and observed distinct AFE usage within the same gene (Fig. 2d, right), with AFEs displaying reciprocal expression across cell types, highlighting cell-type–specific promoter activity and isoform-level regulation.

To further assess FULL-seq’s performance, we analyzed genes with three or more expressed AFEs. FULL-seq successfully identified genes with three (Extended Data Fig. 4c) and four (Extended Data Fig. 4d) distinct AFEs, each validated by junction-spanning reads. In the three AFE example, two AFEs showed similar expression levels, while the third was differentially expressed between conditions, demonstrating complex isoform dynamics that would be missed by gene-level analyses. Importantly, even lowly expressed AFEs showed clear splicing patterns, underscoring FULL-seq’s sensitivity in capturing rare isoforms.

Beyond improving data quality, FULL-seq enhances workflow efficiency, reducing total processing time by approximately 35.1% (∼300 minutes shorter) compared to 10x multi-omics methods. It also avoids the complex, device-dependent protocols that limit scalability in methods such as VASA-seq (Supplementary Fig. 2).

Together, these analyses demonstrate that FULL-seq substantially improves transcript structure resolution in single cells and enables systematic identification of alternative first exons.

### Dissecting AFE usage and promoter complexity through chromatin-transcriptome coupling

The simultaneous measurement of chromatin accessibility and full-length transcript coverage enabled us to investigate the regulatory architecture underlying alternative first exon (AFE) usage, which cannot be resolved by conventional approaches such as 10x Multiome because these methods lack isoform-level resolution (Fig. 3a). Integrated genome browser views revealed clear alignment between ATAC peaks and the transcription start sites (TSSs) of individual AFEs, allowing direct identification of promoter-AFE pairs.

**Fig. 3:**
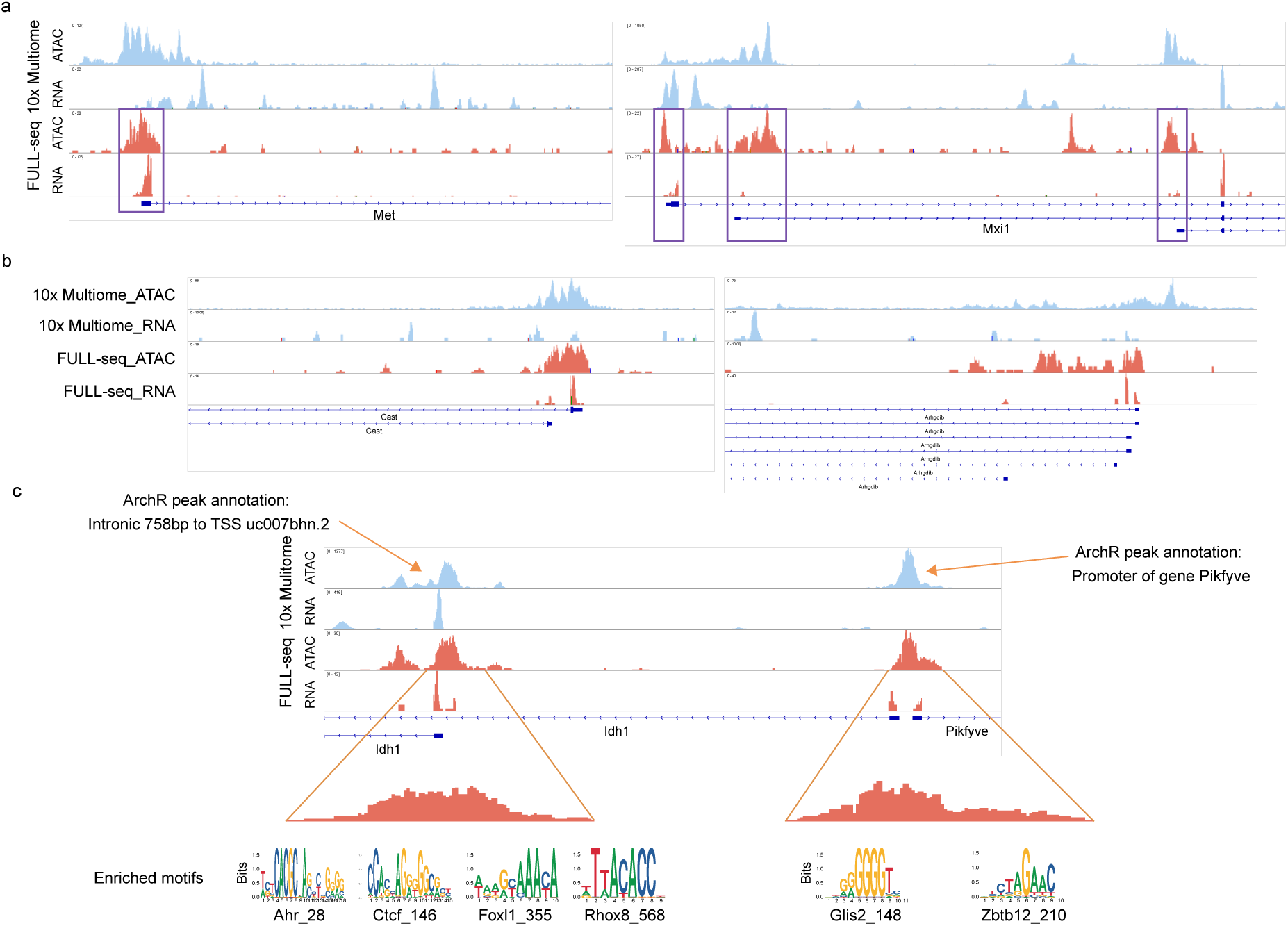
Integrated analysis of chromatin accessibility and AFE usage in CT26 and B16F10 cells using FULL-seq. **a,** IGV genome browser tracks displaying aggregated RNA and ATAC signals for mono-AFE (left) and multi-AFE (three AFEs; right) genes, comparing FULL-seq and 10x Multiome-seq. FULL-seq provides clearer AFE–promoter alignment and more complete transcript coverage. **b,** IGV tracks visualizing chromatin accessibility and AFE expression for genes with multi-AFEs linked to a single ATAC peak. Despite sharing the same accessible region, the AFEs display distinct expression levels, further highlighting that AFE usage does not scale linearly with promoter accessibility. **c,** Integrative view of AFE expression, chromatin accessibility, and motif enrichment at AFE-associated promoter regions. Top panel shows aggregated promoter and AFE signals; middle panel zooms in on promoter peaks with localized ATAC enrichment; bottom panel presents enriched transcription factor motifs, including Ahr_28, which may regulate AFE usage.

In genes with a mono-AFE, we observed strong spatial concordance between promoter ATAC peaks and 5′ RNA signals, indicating a direct relationship between local chromatin accessibility and transcription initiation (Fig. 3a, left). In contrast, genes with multiple AFEs exhibited more complex promoter architectures. For example, the gene *Mxi1*, which expresses three AFEs, displayed three well-aligned ATAC peaks corresponding to distinct 5′ RNA transcription signals (Fig. 3a, right). Compared with 10x Multiome-seq, FULL-seq provided more complete transcript coverage and resolved multiple AFEs within individual genes with greater clarity.

Interestingly, major and minor AFEs within the same gene frequently showed similar levels of promoter accessibility despite markedly different expression levels (Fig. 3a, right; Extended Data Fig. 5a). This observation suggests that chromatin accessibility alone does not scale linearly with AFE usage.

We further identified a spectrum of regulatory topologies illustrating how chromatin accessibility interacts with AFE regulation (Fig. 3b; Extended Data Fig. 5b). In some loci, neighboring genes shared overlapping ATAC peaks yet displayed markedly different transcriptional outputs, with one gene highly expressed and the other weakly expressed or silent. In other cases, multiple AFEs within a single gene were associated with a common accessible promoter region but showed distinct expression levels. These observations suggest the presence of additional regulatory mechanisms, such as coordinated promoter usage or promoter competition, that modulate isoform-specific transcription. Together, these findings demonstrate that FULL-seq resolves complex promoter usage architectures that are obscured by conventional gene-level analyses.

### Identification of Candidate Transcription Factors Regulating AFE Usage

Building on the discovery of diverse promoter architectures, we next sought to identify transcription factors that may regulate AFE usage through motif enrichment analysis of accessible chromatin regions. By analyzing transcription factor (TF) motifs enriched at peaks with differential accessibility between B16F10 and CT26 cells, we identified distinct lineage-associated regulatory signatures (Extended Data Fig. 6a), and ChromVAR revealed cell-to-cell variation in motif accessibility (Extended Data Fig. 6b).

We then examined transcription factor motifs specifically at AFE-associated promoter regions. Genome browser views showed sharp ATAC peaks near TSSs, with distinct motif sets observed at different promoters within the same gene. For example, *Glis2* and *Zbtb12* were identified at the upstream promoter, whereas motifs for *Ahr*, *Ctcf*, *Foxl1*, and *Rhox8* appeared at the downstream promoter (Fig. 3c). These results indicate that individual AFEs within the same gene can be regulated by different transcription factors. Consistent with this interpretation, UMAP projections of motif deviation scores and gene activity revealed coordinated transcription factor activity and expression patterns across cells (Extended Data Fig. 6c).

Notably, conventional gene-centric annotation frameworks such as ArchR^13^ may classify ATAC peaks associated with alternative first-exon promoters as intronic or assign them to neighboring genes. To investigate this limitation, we quantified the conventional genomic annotations of ATAC peaks associated with AFE regions across non-AFE, mono-AFE, and multi-AFE genes (Extended Data Fig. 6d). Only approximately half of these peaks were classified as promoters in non-AFE and mono-AFE genes, with the proportion decreasing to 32.4% in multi-AFE genes. These findings highlight the limitations of gene-centric peak annotations in capturing the regulatory context of alternative first exons.

### FULL-seq profiling in human cancer cell lines

To assess the capability of FULL-seq in human samples, we applied the method to nine human cancer cell lines, including colorectal (HCT116, LS180, KM12, HCT15, HT29, SW620, LS513) and gastric (MKN45, GCIY) lines, yielding 6,257 high-quality single cells after filtering. These cell lines encompass both primary and metastatic origins. Joint analysis of transcriptome and ATAC modalities revealed distinct clustering of the nine cancer cell lines (Fig. 4a), and dot plot visualization confirmed expression of canonical marker genes across all lines (Extended Data Fig. 7a). IGV browser tracks further demonstrated full-length transcript coverage, including long transcripts such as BIRC6 (∼15 kb) and key cell marker genes (Fig. 4b, Extended Data Fig. 7b). Notably, even genes with extremely low expression exhibited read coverage that aligned well with annotated exons (Extended Data Fig. 7b), highlighting the method’s high sensitivity and exon-level resolution.

**Fig. 4:**
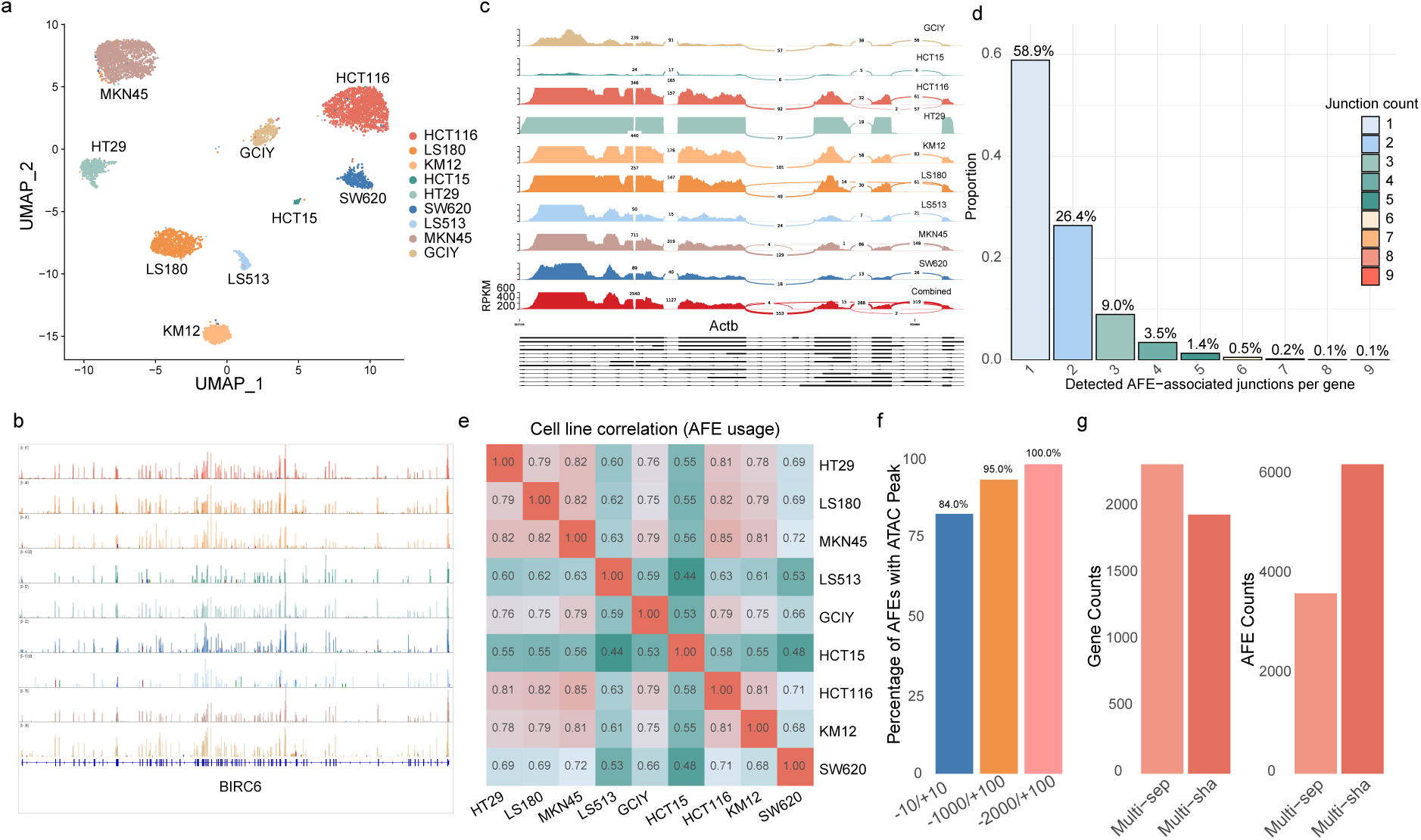
FULL-seq enables isoform-resolved transcriptomics and precise AFE– promoter mapping across nine human cancer cell lines. **a,** UMAP visualization of FULL-seq profiles from nine human cancer cell lines. Cells are clustered using integrated RNA expression and chromatin accessibility data, and cell types are annotated based on the expression of established marker genes. **b,** IGV genome browser tracks showing aggregated full-length mRNA coverage (∼15 kb transcripts) for representative genes across the nine cell types. Even for cell types with relatively few detected reads, transcripts still align cleanly to annotated exons, demonstrating FULL-seq’s sensitivity and ability to capture full-length coverage. **c,** Sashimi plots illustrating AFE usage across the nine cancer cell lines. Exon connectivity is clearly visualized—even with low read depth—highlighting FULL-seq’s ability to resolve exon linkage and isoform structures. Read density is shown as bar heights, and splice junction usage is indicated by arcs, with read counts labeled. **d,** Distribution of the number of distinct detected AFE-associated junctions per gene in the pooled nine-cell-line dataset. Among the genes with at least one detected AFE-associated junction, 58.9% exhibited a single junction and 41.1% exhibited multiple junctions. **e,** Heatmap of AFE expression correlations across nine cancer cell lines, based on FULL-seq RNA data, highlighting cell type–specific transcriptional programs. **f,** Overlap between annotated AFE transcription start sites (TSSs) and ATAC peaks across three genomic windows relative to each TSS (−10 to +10 bp, −1,000 to +100 bp, and −2,000 to +100 bp). Left, proportion of AFEs overlapping at least one ATAC peak. Right, numbers of associated genes and AFEs. **g,** Classification of multi-AFE genes according to promoter-sharing architecture. Although a larger fraction of genes exhibit multi-separate (multi-sep) AFE architectures, multi-shared (multi-sha) AFE genes contain more AFEs per gene, resulting in a substantial fraction of AFEs arising from shared-promoter architectures.

To assess transcript structure detection, we generated Sashimi plots across all nine cell lines, revealing clear exon coverage and well-supported splice junctions, including in cells with relatively low sequencing depth (Fig. 4c). Annotation-guided analysis of the pooled nine-cell-line dataset revealed that 58.9% of genes with detectable AFE-associated junctions exhibited a single distinct junction, whereas 41.1% exhibited multiple junctions (Fig. 4d). Similar distributions were observed across individual cell lines (Supplementary Fig. 3a), highlighting the diversity of AFE-associated transcript structures captured by FULL-seq.

To characterize the conservation and cell-type specificity of AFE-associated genes, we performed intersection analysis across nine cell lines. Multi-AFE genes exhibited broader representation across cell lines, with approximately 72% detected in at least five cell lines, whereas only 28% were detected in fewer than five. In contrast, mono-AFE genes showed more restricted distributions, with approximately 57% detected in fewer than five cell lines and 45% detected in only one or two cell lines (Supplementary Fig. 3b). These findings reveal distinct patterns of AFE-associated gene expression across cancer cell types.

Parallel ATAC-seq data from the same cells confirmed high-quality chromatin accessibility, demonstrated by expected fragment periodicity and strong enrichment at TSSs (Extended Data Fig. 8a–b). Differential analysis revealed distinct chromatin landscapes across tumor types (Extended Data Fig. 8c), supporting FULL-seq’s ability to jointly profile transcriptome and accessibility at single-cell resolution.

### FULL-seq enables precise coupling between chromatin accessibility and isoform-specific promoter usage

We next quantified the spatial alignment between AFE transcription start sites and chromatin accessibility. Genome-wide analysis revealed that approximately 84% of analyzed AFE TSS regions (−10 to +10 bp) overlapped at least one ATAC peak (Fig. 4f). Separately, conventional ArchR annotation classified only 29.9%, 27.4%, and 23.6% of AFE-associated peaks as promoters in non-AFE, mono-AFE, and multi-AFE genes, respectively (Extended Data Fig. 8d). Together, these findings illustrate the close spatial association between AFE transcription initiation and accessible chromatin, while highlighting the limitations of conventional gene-centric peak annotations in capturing alternative promoter contexts. Representative IGV views further illustrated that broader TSS-associated regions encompass additional nearby ATAC peaks (Extended Data Fig. 8e).

We further categorized multi-AFE genes into two types: multi-separate (multi-sep) AFE genes, where each AFE is associated with a distinct promoter, and multi-shared (multi-sha) AFE genes, where multiple AFEs share a common promoter. More genes were classified as multi-sep AFE, though multi-sha AFE genes tended to have a greater number of AFE isoforms (Fig. 4g).

Together, these findings demonstrate that chromatin accessibility and transcription factor activity can be linked directly to promoter usage and full-length transcript isoforms. By resolving regulatory relationships at the promoter–isoform level, FULL-seq distinguishes isoform-specific regulatory programs that are obscured by conventional gene-level analyses (Extended Data Fig. 1a). Thus, FULL-seq moves beyond gene-level chromatin– expression associations to map regulatory events at the promoter–isoform interface.

### AFE usage reveals functionally relevant isoform programs associated with cancer metastasis

To investigate whether AFE usage captures cancer-state differences that are not apparent at the gene level, we compared AFE expression between primary and metastatic colorectal cancer cell lines. Exploratory analysis of AFE-associated junction expression revealed distinct patterns among the analyzed colorectal cancer cell lines (Fig. 5a). AFE-level profiling revealed candidate expression differences between the analyzed colorectal cancer cell lines that were not readily apparent from aggregate gene-expression patterns (Extended Data Fig. 9a). Affected genes include well-characterized regulators of metastatic processes, including apoptosis (*BAG1*^19,20^), metabolic rewiring (*LDHA*^21,22^, *SLC25A5*^23^), cytoskeletal organization (*TPM1*^24,25^), RNA processing (*HNRNPU*^26,27^), and stress response signaling (*XBP1*^28,29^). While these genes are known to contribute to cancer progression, the contribution of AFE usage to cancer progression remains poorly understood.

**Fig. 5:**
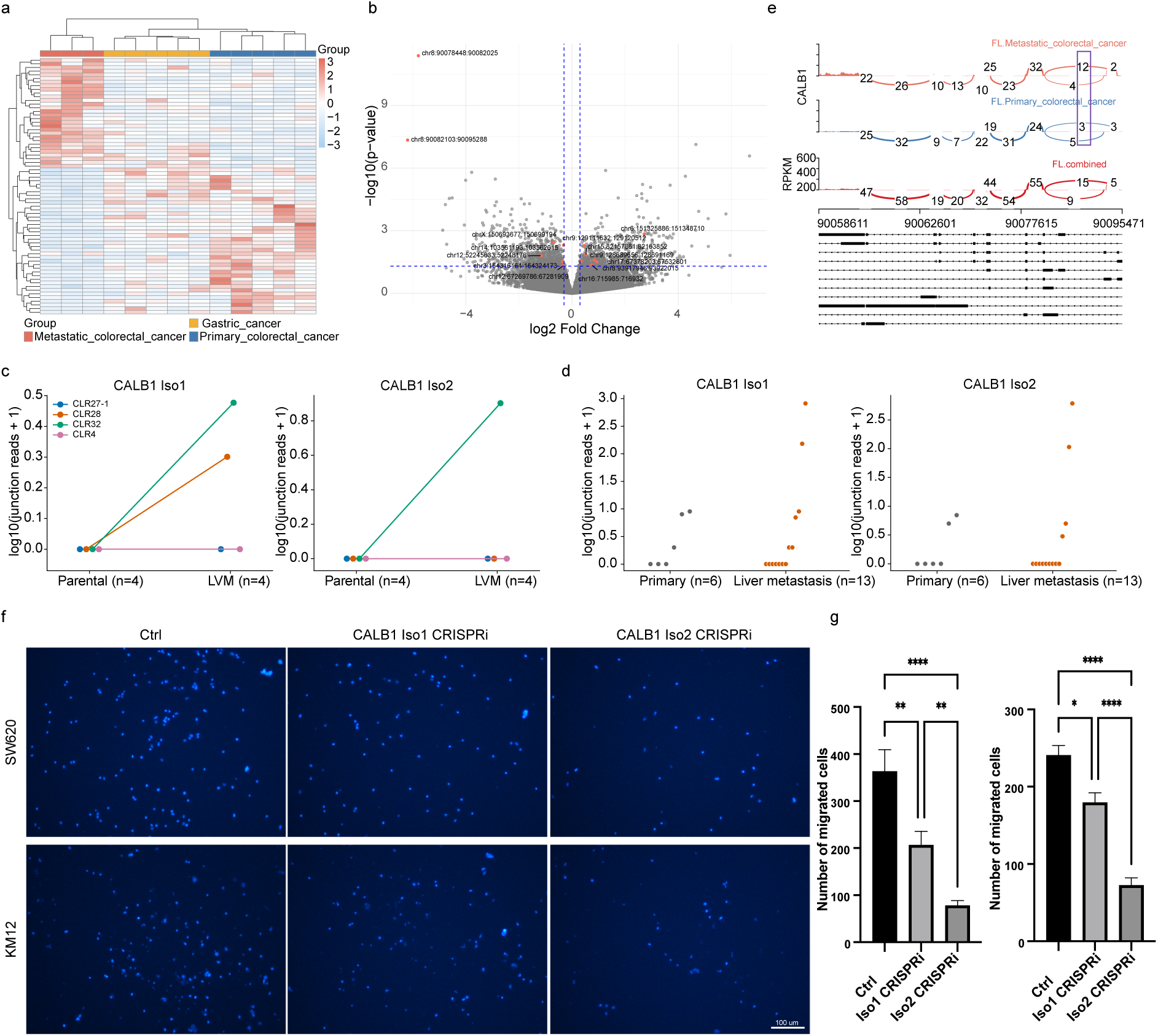
AFE usage reveals functionally relevant isoform programs associated with colorectal cancer metastasis. **a**, Heatmap of normalized AFE expression (z-scores) across gastric, primary colorectal, and metastatic colorectal cancer cell lines, showing distinct subgroup-specific AFE usage profiles. **b,** Volcano plot of differential AFE usage in colorectal cancer cell lines from the Cancer Cell Line Encyclopedia (CCLE), comparing cell lines derived from primary tumors (n = 21) and metastatic sites (n = 17). Red points indicate AFE events also identified as candidate metastasis-associated AFEs in the FULL-seq analysis. **c,** CALB1 Iso1- and Iso2-specific junction abundance in matched parental and liver-metastatic (LVM) derivatives from the GSE138248 PDX model (n = 4 pairs). Matched samples are connected by lines. **d,** CALB1 Iso1- and Iso2-specific junction abundance in primary colorectal tumors (n = 6) and liver metastases (n = 13) from the GSE221240 patient cohort. Individual samples are shown as points. Values in **c,d** are log10(junction reads + 1). **e**, Sashimi plots showing differential CALB1 alternative first exon usage across gastric, primary colorectal, and metastatic colorectal cancer cell lines. Read coverage is represented by bar heights, and splice junction usage by arcs, with read counts labeled. Red boxes highlight the metastasis-associated CALB1 AFEs. **f,** Representative Transwell migration images of SW620 and KM12 colorectal cancer cells following CRISPRi-mediated repression of CALB1 Iso1 or Iso2 compared with control cells. Nuclei were visualized by DAPI staining. g, Quantification of migrated SW620 (left) and KM12 (right) cells following CRISPRi-mediated repression of CALB1 Iso1 or Iso2 using a pool of three sgRNAs targeting each isoform. Statistical significance was determined by one-way ANOVA followed by Dunnett’s multiple-comparisons test, with each CALB1 isoform knockdown compared with the control. * P < 0.05, **P < 0.01, ***P < 0.001, ****P < 0.0001.

To independently evaluate metastasis-associated AFE usage, we analyzed 38 colorectal cancer cell lines from the Cancer Cell Line Encyclopedia (CCLE), comprising 21 derived from primary tumors and 17 from metastatic sites. Differential AFE analysis revealed widespread differences between primary and metastatic colorectal cancer cell lines, including a subset of AFE events that overlapped those identified by FULL-seq (Fig. 5b). Among these recurrent events, two AFEs of CALB1—chr8:90078448:90082025 (Iso1) and chr8:90082103:90095288 (Iso2)—with transcription start sites separated by approximately 13.3 kb, were enriched in metastatic cancers in both the FULL-seq and public datasets (Fig. 5b).

We further examined CALB1 AFE expression in a matched parental and liver-metastatic derivative PDX dataset (GSE138248). CALB1 AFE-specific junctions were detected in a subset of liver-metastatic derivatives, although their abundance was generally low and was not accompanied by a consistent increase in total CALB1 expression (Fig. 5c and Extended Data Fig. 9b). In an independent patient cohort (GSE221240), CALB1 AFE expression was heterogeneous, with marked activation observed in a subset of liver metastases (Fig. 5d). Together, these analyses provide complementary evidence for metastasis-associated CALB1 AFE activation across colorectal cancer models and patient tumors, while highlighting its heterogeneous occurrence.

We next examined *CALB1* as a representative metastasis-associated AFE event. Sashimi plot analysis resolved distinct *CALB1* transcripts initiated from alternative first exons and revealed differential usage of these AFEs between primary and metastatic colorectal cancer cells (Fig. 5e). In particular, the metastasis-associated *CALB1* AFEs showed increased usage in metastatic cells, whereas this transcript-level difference was not readily apparent from aggregate gene-level measurements. Thus, *CALB1* illustrates how alternative transcription initiation can generate cancer-state-specific transcript outputs that are masked by conventional gene-level analysis.

### CRISPRi targeting of CALB1 AFEs alters colorectal cancer cell migration

To investigate the functional relevance of the metastasis-associated CALB1 AFE, we employed CRISPR interference (CRISPRi) to suppress its expression. Three independent sgRNAs were designed to target the promoter region associated with this AFE. RT–qPCR confirmed effective suppression of the targeted CALB1 AFE following CRISPRi (Extended Data Fig. 9c).

We next assessed the functional consequence of *CALB1* AFE suppression using Transwell migration assays in two colorectal cancer cell lines. CRISPRi-mediated repression of the metastasis-associated *CALB1* AFEs reduced cell migration compared with control cells, as observed in representative Transwell images (Fig. 5f). Quantification confirmed a consistent reduction in migratory capacity following *CALB1* AFE suppression in both cell lines (Fig. 5g). Analysis of individual sgRNAs provided additional support for the observed phenotype (Extended Data Fig. 9d–g).

Together, these results demonstrate that AFE-resolved analysis can uncover functionally relevant transcript programs that are invisible at the gene level. The identification, independent validation, and CRISPRi-mediated perturbation of metastasis-associated *CALB1* AFEs establish alternative promoter and first-exon usage as a biologically consequential layer of cancer-state regulation and illustrate how FULL-seq can connect isoform-resolved regulatory architecture to cellular phenotype.

## Discussion

Single-cell multiomic technologies have enabled joint profiling of chromatin accessibility and gene expression, but their interpretation remains largely gene-centric. FULL-seq addresses this limitation by jointly capturing chromatin accessibility and full-length transcript coverage from the same cell, enabling regulatory relationships to be resolved at the promoter–isoform level.

Using FULL-seq, we uncover diverse relationships between promoter accessibility and AFE expression. Genes exhibit distinct promoter–AFE architectures, including similarly accessible promoters associated with unequal transcript outputs and multiple independent promoters generating distinct isoforms. Moreover, alternative promoters within the same gene can exhibit distinct transcription factor regulatory programs, highlighting regulatory heterogeneity at the isoform level. Together, these findings demonstrate how gene-level frameworks can obscure or misassign regulatory relationships in multi-promoter genes.

Importantly, AFE-resolved analysis revealed metastasis-associated transcript programs that were not apparent at the gene-expression level and were reproduced in independent colorectal cancer datasets. CRISPRi-mediated repression of metastasis-associated *CALB1* AFEs reduced colorectal cancer cell migration, providing functional evidence that isoform-specific regulation can contribute to cancer-associated phenotypes. Together, our findings establish the promoter–isoform interface as a functionally relevant dimension of gene regulation and position FULL-seq as a framework for connecting chromatin accessibility, transcript identity, and cellular phenotype beyond the gene level.

## Method

### Cell culture

MKN45, HCT15, KM12, LS513, and CT26 cells were cultured in RPMI 1640 medium (11875-093, ThermoFisher) supplemented with 10% FBS (16000044, ThermoFisher) and 1% penicillin-streptomycin (15140122, ThermoFisher). HT29, SW620, HCT116, and B16F10 cells were cultured in Dulbecco’s Modified Eagle Medium (DMEM, 11965092, ThermoFisher) with the addition of 10% FBS and 1% of penicillin-streptomycin. GCIY and LS180 cells were cultured in DMEM/F-12 Medium (D/F12, 21331020, ThermoFisher) with the addition of 10% FBS and 1% of penicillin-streptomycin. Cells were incubated at 37 °C in 5% CO2 and maintained at the exponential phase. All cells were digested with Trypsin-EDTA (0.25%) (25200114, ThermoFisher) for preparing single-cell suspension.

### Investigate reverse transcription using biotinylated dNTP

To evaluate the efficiency and feasibility of reverse transcription with biotinylated nucleotides, we tested three types of biotinylated dNTPs: biotinylated dUTP (BU), biotinylated dATP (BA), and biotinylated dCTP (BC). GAPDH mRNA was used as the template for all reactions. Each reverse transcription (RT) reaction contained 2.5 μl of 10 mM dNTP mix, 2.5 μl of 1 mM biotinylated dUTP/dATP/dCTP, 2.5 μl of Maxima H Minus Reverse Transcriptase (Thermo Fisher Scientific, Cat. No. EP0753), 1 μl of SUPERase In RNase inhibitor, 2 μl of GAPDH mRNA (50 ng/μl), 2.5 μl of 100 μM oligo(dT)₁₈ primer, 10 μl of 5× RT buffer, and 27 μl of nuclease-free water. Reverse transcription was carried out at 50 °C for 30 min, followed by enzyme inactivation at 85 °C for 5 min and a hold at 10 °C.

RT products were purified using 1.8× SPRIselect beads (Beckman Coulter, Cat. No. B23317) according to the manufacturer’s protocol. Briefly, 90 μl of resuspended beads (1.8×) were added to each reaction, mixed by pipetting, and incubated for 5 min at room temperature. The samples were placed on a magnetic rack until the solution cleared, and the supernatant was removed. Beads were washed twice with 80% ethanol (300 μl and 200 μl, respectively), air-dried, and eluted in 10.5 μl of nuclease-free water. After a 2 min incubation at room temperature, eluates were briefly centrifuged, placed on a magnet, and the supernatants were transferred to fresh tubes. Purified RNA/cDNA hybrid was stored at 4 °C for up to 72 h or at −20 °C for long-term storage. Fragment size and concentration were assessed using the Agilent Bioanalyzer High Sensitivity DNA chip.

### Optimization of reverse transcription efficiency using biotinylated dCTP

To further optimize reverse transcription efficiency, we evaluated the incorporation performance of biotinylated dCTP at varying molar ratios to unmodified dCTP (biotinylated dCTP:dCTP = 0:10, 2:8, 4:6, 5:5, 6:4, 10:0). GAPDH mRNA was used as the template for all reactions. Each reverse transcription (RT) reaction contained 2.5 μl of 10 mM dATP + dTTP + dGTP mix, 2.5 μl of 1 mM biotinylated dCTP + dCTP mix at the indicated ratios, 2.5 μl of Maxima H Minus Reverse Transcriptase (Thermo Fisher Scientific, Cat. No. EP0753), 1 μl of SUPERase In RNase inhibitor, 2 μl of GAPDH mRNA (50 ng/μl), 2.5 μl of 100 μM oligo(dT)₁₈ primer, 10 μl of 5× RT buffer, and 27 μl of nuclease-free water. Reverse transcription was carried out at 50 °C for 30 min, followed by enzyme inactivation at 85 °C for 5 min and a hold at 10 °C.

RT products were purified using 1.8× SPRIselect beads (Beckman Coulter, Cat. No. B23317) according to the manufacturer’s protocol. Briefly, 90 μl of resuspended beads (1.8×) were added to each reaction, mixed by pipetting, and incubated for 5 min at room temperature. The samples were placed on a magnetic rack until the solution cleared, and the supernatant was removed. Beads were washed twice with 80% ethanol (300 μl and 200 μl, respectively), air-dried, and eluted in 10.5 μl of nuclease-free water. After a 2 min incubation at room temperature, eluates were briefly centrifuged, placed on a magnet, and the supernatants were transferred to fresh tubes. Purified RNA/cDNA hybrid was stored at 4 °C for up to 72 h or at −20 °C for long-term storage. Fragment size and concentration were assessed using the Agilent Bioanalyzer High Sensitivity DNA chip.

### FULL-seq

#### Nuclei preparation

All procedures were performed on ice unless otherwise indicated. To minimize sample loss, centrifugations were carried out at 4 °C using a swing-bucket rotor, and care was taken to avoid disturbing the pellet when removing supernatants. Before processing, 1.5 ml tubes were coated with 0.5 ml of 0.5× DPBS containing 0.5% (w/v) BSA and RNase inhibitor (RI, 0.2 U/μl) by inverting the tube three to five times, followed by removal of all liquid from the tube and cap. Coating the tubes reduces cell adhesion and is particularly important when handling low cell numbers. Cells were counted using a C-Chip disposable hemocytometer, and viability was confirmed to exceed 90% by Trypan Blue staining. Approximately 1 × 10⁵ cells per sample were pelleted by centrifugation at 500 × *g* for 5 min, washed once with 500 μl ice-cold 0.5× DPBS containing 0.5% BSA and RI, and centrifuged again under the same conditions. The pellet was then resuspended in 50 μl ice-cold RSB-DTN-RI buffer (Mix 487.5 μl Omni-ATAC RSB, 5 μl 10% Tween-20, 5 μl 10% IGEPAL CA-630 and 2.5 μl 2% Digitonin, 1.25ul RNase inhibitor (400x). Make fresh) and incubated on ice for 3 min to lyse the cytoplasmic membrane. Subsequently, 1 ml of ice-cold RSB-T-RI buffer (Mix 10 ml Omni-ATAC RSB and 100 μl 10% Tween-20, 25ul RNase inhibitor (400x)) was added, and the suspension was gently mixed by inversion. The resulting nuclei were counted and collected by centrifugation at 1,000 × *g* for 8 min at 4 °C. After carefully removing the supernatant, the purified nuclei were kept on ice and resuspended in the reverse transcription (RT) mix immediately before the next step.

#### Reverse transcription

Following nuclei isolation, nuclei were pelleted and washed twice with ice-cold 0.5× DPBS containing 0.5% (w/v) BSA and RiboLock RNase inhibitor. Approximately 1 × 10⁵ nuclei were resuspended in 50 μl of reverse transcription (RT) mix containing 2.5 ul 100uM oligo_dT, 2.5 ul 10 mM dATP/dTTP/dGTP, 1.5 ul 10 mM Bio-dCTP, 1 ul 10 mM dCTP, 3.5ul Maxima H Minus Reverse Transcriptase (Thermo Fisher Scientific, Cat. No. EP0753), 1 ul 40 U/μl RiboLock RNase Inhibitor, 1ul 20 U/μl SUPERaseIn RNase Inhibitor, 10 ul 5× RT Buffer, 12 ul 50% PEG6000, 15 ul nuclease-free water. The RT reaction was transferred to a new PCR tube and incubated in a thermocycler at 50 °C for 10 min, followed by three thermal cycles of 8 °C for 12 s, 15 °C for 45 s, 20 °C for 45 s, 30 °C for 30 s, 42 °C for 2 min, and 50 °C for 3 min, with a final extension at 50 °C for 5 min. After completion, the RT mixture was transferred to a pre-coated, chilled 1.5 ml tube containing 1 ml of ice-cold 0.5× DPBS with 0.5% BSA, and centrifuged at 1,000 × g for 5 min at 4 °C. The pellet was washed once with 0.5 ml of diluted nuclei buffer and centrifuged again under the same conditions. The resulting nuclei were resuspended in 20 μl of diluted nuclei buffer (50ul 20x Nuclei Buffer (PN-2000207, 10x Genomics), 1 ul 1M DTT, 25 ul RNase Inhibitor, 924 ul nuclease-free water), yielding a concentration of approximately 10,000 nuclei/μl. The nuclei concentration was determined using a hemocytometer and adjusted to 5,000 nuclei/μl with diluted nuclei buffer for subsequent processing.

#### RNA–DNA hybrid and DNA tagmentation

Following reverse transcription, nuclei containing RNA–DNA hybrids were subjected to tagmentation to fragment genomic DNA and RNA–DNA hybrids, then incorporate sequencing adapters. The transposition reaction was prepared on ice by combining 7 μl of ATAC Buffer B (10x Genomics, Cat. No. 2000193) and 3 μl of ATAC Enzyme B (10x Genomics, Cat. No. 2000265). The mixture was pipette-mixed ten times, briefly centrifuged, and 10 μl of the Transposition Mix was aliquoted into PCR strip tubes, one per sample. The appropriate volumes of nuclei stock and diluted nuclei buffer were calculated according to the Nuclei Concentration Guidelines to achieve a final volume of 5 μl. The corresponding amount of diluted nuclei buffer was first added to the Transposition Mix, followed by gentle mixing and brief centrifugation. The nuclei suspension was then gently pipette-mixed six times (pipette set to 10 μl) and added directly to the Transposition Mix. Samples were not centrifuged after mixing. The transposition reaction was carried out in a thermocycler with a heated lid set to 50 °C, using the following conditions: 37 °C for 60 min, followed by a hold at 4 °C.

#### GEM Generation & Barcoding

After tagmentation, GEM were generated using the 10x Chromium NextGEM Multiome kit (CG000338) following the manufacturer’s recommended protocol. Briefly, following tagmentation, nuclei were encapsulated and barcoded using the Chromium Next GEM Single Cell Multiome ATAC + Gene Expression system (10x Genomics). The barcoding master mix was prepared on ice by combining Barcoding Reagent Mix (Cat. No. 2000267), Reducing Agent B (Cat. No. 2000087), and Barcoding Enzyme Mix (Cat. No. 2000266/2000273) in the order listed, for a total reaction volume of 60 μl per sample. The master mix was pipette-mixed ten times and briefly centrifuged before use.

For each reaction, 60 μl of the master mix was added to 15 μl of transposed nuclei, yielding a total of 75 μl. The mixture was gently pipette-mixed and loaded into row 1 of a Chromium Next GEM Chip J, while Gel Beads and Partitioning Oil were loaded into rows 2 and 3, respectively, according to the manufacturer’s protocol. Unused wells were filled with 50–150 μl of 50% glycerol solution to maintain chip hydration. The loaded chip was immediately placed into the Chromium Controller (10x Genomics) for GEM generation (∼18 min).

After completion, GEMs were carefully recovered from the wells, ensuring that samples appeared opaque and uniform, and transferred into chilled 8-tube PCR strips on ice. The GEMs were then incubated in a thermal cycler (Bio-Rad C1000 Touch, 125 μl reaction volume, 50 °C lid temperature) with the following program: 37 °C for 45 min, 25 °C for 30 min, and a 4 °C hold. Reactions were quenched by adding 5 μl Quenching Agent to each sample, followed by gentle mixing (10×). The quenched GEMs were stored at −80 °C for up to four weeks or immediately processed for downstream clean-up and amplification.

#### Post-GEM incubation cleanup

After GEM incubation, reactions were cleaned to remove oil, barcoding reagents, and residual enzymes. Cleanup was performed sequentially using Dynabeads MyOne SILANE and SPRIselect reagents (Beckman Coulter) according to the manufacturer’s protocol with minor modifications.

For the Dynabeads cleanup, 125 μl Recovery Agent (10x Genomics, Cat. No. 220016) was added to each sample at room temperature, mixed gently by inversion, and briefly centrifuged to separate the biphasic layers. The lower pink layer containing Recovery Agent and Partitioning Oil was carefully removed without disturbing the aqueous phase. A Dynabeads Cleanup Mix consisting of Cleanup Buffer (Cat. No. 2000088), Dynabeads MyOne SILANE (Cat. No. 2000048), and Reducing Agent B (Cat. No. 2000087) was freshly prepared, and 200 μl of this mix was added to each sample. Samples were mixed by pipetting ten times and incubated for 10 min at room temperature. Beads were then captured on a magnetic separator (high position), washed twice with 80% ethanol, air-dried, and eluted in 50.5 μl of Elution Solution I containing Buffer EB, 10% Tween-20, and Reducing Agent B. The eluate was mixed gently, incubated for 1 min at room temperature, and transferred to a new tube after magnetic separation.

For the SPRIselect cleanup, 90 μl of fully resuspended SPRIselect beads were added to each sample and mixed thoroughly. After a 5 min room-temperature incubation, the beads were magnetically separated, and the supernatant was removed. The beads were washed twice with 200 μl of freshly prepared 80% ethanol, air-dried, and eluted in 46.5 μl of Buffer EB. Samples were mixed gently, incubated for 2 min at room temperature, centrifuged briefly, and placed on the magnet until the solution cleared. A total of 46 μl of cleaned product was transferred to a new tube and stored at −20 °C until use.

#### Separation and purification of DNA and RNA

To separate biotinylated RNA/DNA hybrids and genomic DNA fractions, streptavidin bead–based capture was performed using Dynabeads MyOne C1 (65001, Thermo Fisher Scientific). Beads were divided into two groups (A and B) and pre-washed to remove preservatives and equilibrate binding conditions. Specifically, 20 μl (group A) and 10 μl (group B) of Dynabeads were transferred into separate 1.5 ml microcentrifuge tubes and washed three times with 500 μl of 1× B&W-T buffer (5 mM Tris-HCl pH 7.5, 0.5 mM EDTA, 1 M NaCl, 0.01% Tween-20), followed by one wash with 500 μl of 1× B&W-T buffer containing RNase inhibitor (B&W-T/RI). Beads were placed on a magnetic rack for ∼1 min between washes to allow the solution to clear before removing the supernatant. After the final wash, group A beads were resuspended in 50 μl of 2× B&W buffer, while group B beads were kept on ice until use.

Each sample was incubated with 100 μl of group A beads on an end-to-end rotator at room temperature (RT) for 60 min at 40 rpm to capture biotin-labeled cDNA/RNA hybrids. Beads were magnetically separated, and the supernatant was transferred to the group B beads for a second 60 min rotation under the same conditions. The resulting supernatant, containing chromatin and unbound DNA fragments, was collected for downstream library preparation.

The beads from groups A and B were then pooled and washed with 200 μl of 1× B&W-T/RI buffer twice, and all supernatant were added to the previously collected chromatin fraction. The cDNA/RNA-bound beads were washed three additional times with 500 μl of 1× B&W-T/RI buffer at RT without resuspending, and the washed beads were stored at RT for short-term stability or at 4 °C for extended storage prior to elution or downstream processing.

#### Final RNA library construction

RNA libraries were constructed from bead-bound cDNA/RNA hybrids following streptavidin capture. For gap filling, MyOne C1 beads carrying cDNA/RNA hybrids were resuspended in 81 μl of gap-filling buffer containing 2 μl Maxima H Minus Reverse Transcriptase (Thermo Fisher Scientific), 41 μl 2× NEBNext Mix, and 38 μl nuclease-free water. The reaction was incubated at 50 °C for 15 min and 72 °C for 5 min to fill the ∼9-bp gaps generated during Tn5 transposition, followed by enzyme inactivation at 85 °C for 5 min.

For library amplification and indexing, PCR was performed using 7.5 μl SI-PCR Primer B (P5), 2.5 μl sample-specific Index N (P7), and 9 μl 2× NEBNext Mix. The PCR mixture was added to the bead suspension and mixed thoroughly. Thermal cycling was carried out as follows: 72 °C for 5 min; 98 °C for 30 s; 15 cycles of 98 °C for 10 s, 67 °C for 30 s, and 72 °C for 30 s; a final extension at 72 °C for 5 min; and a hold at 4 °C.

A 5 μl aliquot of PCR product was purified and quantified to determine the optimal cycle number for each sample, preventing over-amplification.

Post-PCR cleanup and size selection were performed using a double-sided SPRIselect purification. Briefly, 60 μl of well-resuspended SPRIselect reagent (0.6×) was added to each reaction, mixed, and incubated for 5 min at room temperature. The mixture was placed on a magnetic rack (high position) until clear, and 150 μl of supernatant was transferred to a new tube. A second round of bead binding was performed by adding 95 μl SPRIselect reagent (1.55×) to the recovered supernatant, followed by 5 min incubation at room temperature. Beads were captured magnetically, washed twice with 80% ethanol (300 μl and 200 μl), briefly centrifuged, and air-dried. DNA was eluted in 20.5 μl Buffer EB, incubated for 2 min at room temperature, and the eluate was cleared on the magnet. A total of 20 μl purified library was transferred to a new tube and stored at 4 °C for up to 72 h or at −20 °C for long-term storage.

#### Final ATAC library construction

ATAC libraries were constructed from 500 ul DNA fragments remaining after RNA/DNA hybrid separation. Following bead purification, samples were prepared for amplification and indexing. SPRIselect reagent (Beckman Coulter) was vortexed until fully resuspended, and 900 μl was added to each sample. The mixture was pipette-mixed thoroughly, incubated for 5 min at room temperature (RT), and centrifuged briefly. Samples were placed on a magnetic rack (high position) until clear, and the supernatant was removed. The beads were washed twice with 500 μl of freshly prepared 80% ethanol, centrifuged briefly, placed on the magnet (low position), and residual ethanol was carefully removed to prevent inhibition of PCR. Beads were immediately resuspended in 40.5 μl Buffer EB, gently mixed, incubated for 2 min at RT, and cleared magnetically. A total of 40 μl of eluate was transferred to a new tube for PCR amplification.

For ATAC library amplification and indexing, 40 μl of eluate was combined with 60 μl ATAC PCR mix containing 50 μl 2× NEBNext Mix, 7.5 μl SI-PCR Primer B (P5), and 2.5 μl individual sample Index N (P7). Reactions were mixed thoroughly and amplified under the following thermal cycling conditions: 72 °C for 5 min; 98 °C for 30 s; 15 cycles of 98 °C for 10 s, 67 °C for 30 s, and 72 °C for 30 s; followed by a final extension at 72 °C for 5 min and a hold at 4 °C.

Post-PCR double-sided size selection was performed using SPRIselect. Specifically, 60 μl of SPRIselect reagent (0.6×) was added to each reaction, mixed, and incubated for 5 min at RT. The mixture was placed on a magnetic rack until clear, and 150 μl of supernatant was transferred to a new tube. A second binding step was performed by adding 95 μl of SPRIselect reagent (1.55×) to the recovered supernatant, followed by 5 min incubation at RT. Beads were captured magnetically, washed twice with 80% ethanol (300 μl and 200 μl), air-dried briefly, and eluted in 20.5 μl Buffer EB. The eluate was incubated for 2 min at RT, cleared magnetically, and 20 μl of purified ATAC library was transferred to a new tube. Libraries were stored at 4 °C for up to 72 h or at −20 °C for long-term preservation.

#### RNA and ATAC libraries preprocessing and quality control

Sequencing reads from both RNA and ATAC libraries were processed using Cell Ranger ARC (10x Genomics). Because FULL-seq RNA libraries were generated through an ATAC-style workflow, the RNA reads carried ATAC barcodes rather than standard RNA barcodes. To ensure compatibility with Cell Ranger, ATAC barcodes were first converted to the corresponding RNA barcode whitelist provided by 10x Genomics (Chromium Next GEM Single Cell Multiome ATAC + Gene Expression). Specifically, the raw FASTQ files were parsed to replace each ATAC barcode sequence in read 1 with its matched RNA barcode based on the whitelist table (737K-arc-v1_whitelist.txt), ensuring consistent barcode structure. After barcode conversion and UMIs added, the modified FASTQ files were processed with the standard Cell Ranger ARC pipeline to generate feature–barcode matrices for both RNA and ATAC modalities.

Low-quality cells with fewer than 1000 (mouse), or 700 (human) detected genes, more than 8% (mouse), 30% (human) mitochondrial reads, or fewer than 300 ATAC features were filtered out. The remaining high-quality cells were analyzed using Seurat v5^30^.

#### Gene-level quantification of AFE-associated junctions

To characterize AFE-associated junction multiplicity, annotated AFE events were identified using SUPPA2 and reference GTF annotations. STAR-derived splice junctions associated with annotated first exons were assigned to their corresponding genes using a gene–junction mapping table. Duplicate gene–junction pairs were removed, and distinct junctions were indexed within each gene. Junction detection was evaluated from the corresponding FULL-seq RNA count matrices, and the number of distinct detected AFE-associated junctions was calculated for each gene. An AFE-associated junction was considered detected if it had a nonzero count in at least one cell within the analyzed dataset. For gene-level distributions, each gene was counted once, and proportions were calculated relative to all genes with at least one detected AFE-associated junction.

Non-AFE genes contained only a single annotated first exon/TSS. Genes with multiple annotated AFEs were further classified as mono-AFE when only one AFE was detectably expressed, or multi-AFE when two or more AFEs were expressed. Multi-AFE genes were further subdivided based on promoter sharing: multi-sep AFE (distinct promoters), in which AFEs were associated with non-overlapping ATAC peaks, and multi-sha AFE (shared promoters), in which AFEs mapped to the same proximal promoter peaks.

To assess the spatial relationship between AFE transcription start sites (TSSs) and chromatin accessibility, three genomic windows were defined relative to each annotated AFE TSS: −10 to +10 bp, −1,000 to +100 bp, and −2,000 to +100 bp, where negative and positive coordinates indicate upstream and downstream positions relative to the transcriptional direction, respectively. These intervals were intersected with ATAC peak regions. An AFE was considered overlapping if at least one ATAC peak intersected its corresponding TSS-associated window. Overlap frequencies were calculated as the proportion of analyzed AFEs with at least one overlapping ATAC peak within each window.

#### Chromatin accessibility and peak calling

ATAC fragments were processed using ArchR for quality control, TSS enrichment scoring, and fragment length periodicity assessment. Peaks were called with MACS3. Promoter accessibility was quantified within ±2 kb of annotated transcription-start sites. Differentially accessible peaks between cell types were identified using ArchR’s “getMarkerFeatures” function with FDR < 0.05. Single-cell motif deviation scores were computed using ChromVAR, and TF-activity UMAPs were generated to compare motif accessibility and TF expression.

#### Transcription-factor motif analysis

Motif enrichment analysis was performed using the **Signac** package in R. Promoter and enhancer ATAC peaks associated with AFE-regulated genes were analyzed separately. Transcription factor binding motifs were identified by scanning these peaks against position weight matrices (PWMs) from the **JASPAR 2022** database using the *FindMotifs* function.

#### Exploratory analysis of cancer-associated AFE expression patterns

Single cells were grouped according to their annotated cell-line origins into gastric, primary colorectal, and metastatic colorectal cancer categories. Equal numbers of cells were randomly sampled from each group and aggregated to generate pseudobulk profiles for exploratory analysis. An initial DESeq2 analysis was used to prioritize candidate AFE-associated junctions (adjusted P < 0.05), and normalized expression values were transformed into z-scores for visualization. The resulting patterns were interpreted descriptively at the cell-line-group level.

#### Isoform-specific repression

To functionally evaluate metastasis-associated AFE usage, isoform-specific CRISPR interference (CRISPRi) was performed by targeting promoter regions associated with alternative *CALB1* first exons. Stable dCas9-expressing SW620 and KM12 colorectal cancer cell lines were generated by lentiviral transduction followed by blasticidin selection. Lentiviral particles were produced using psPAX2 and pMD2.G packaging plasmids together with the dCas9 construct.

Three independent sgRNAs were designed for each *CALB1* isoform to target the promoter/TSS region associated with the corresponding AFE while avoiding regions shared by alternative AFEs. sgRNA constructs were packaged into lentivirus and transduced into dCas9-expressing cells, followed by puromycin selection. Cells transduced with an empty sgRNA vector were used as controls. The three sgRNAs targeting each *CALB1* AFE were pooled for the primary functional analysis, with individual sgRNAs analyzed separately to assess reproducibility. sgRNA sequences are provided in Supplementary Table 1.

#### RT–qPCR validation of *CALB1* AFE repression

The efficiency of CRISPRi-mediated repression was assessed by RT–qPCR using AFE-specific primers. Primer pairs were designed to detect CALB1 Iso1 (NM_004929) and Iso2 (ENST00000523716.5). Expression of the corresponding targeted CALB1 AFE was measured following CRISPRi and normalized to GAPDH. Primer sequences are provided in Supplementary Table 2.

#### Transwell migration assay

Cell migration was assessed using 24-well Transwell inserts containing 8.0-µm-pore polycarbonate membranes (Corning, Cat. No. 3422). SW620 and KM12 cells expressing dCas9 and the indicated *CALB1* AFE-targeting sgRNAs or control sgRNA were harvested and resuspended in serum-free culture medium. Cells were seeded into the upper chamber in 100 µl serum-free medium, and 600 µl medium supplemented with fetal bovine serum was added to the lower chamber as a chemoattractant.

Following incubation, non-migrated cells remaining on the upper surface of the membrane were gently removed with a cotton-tipped applicator. Cells that migrated to the underside of the membrane were fixed with 4% paraformaldehyde for 20 min, washed three times with PBS, and stained with DAPI for 10 min. Migrated cells were imaged using an inverted microscope at ×100 magnification and quantified for statistical analysis.

For comparisons among control and *CALB1* AFE knockdown groups, statistical significance was assessed by one-way ANOVA followed by Dunnett’s multiple-comparisons test versus control.

#### Statistical analysis and visualization

All statistical analyses were conducted in R v4.3. Differential expression and chromatin accessibility at the single-cell level were assessed using non-parametric Wilcoxon rank-sum tests, whereas pseudobulk differential analyses were performed with DESeq2 using the Wald test. Correlations were computed using Pearson’s correlation coefficient unless otherwise specified. Motif enrichment *P* values were corrected using the Benjamini– Hochberg false discovery rate (FDR) method. All plots were generated using ggplot2, ComplexHeatmap, IGV, and Sashimi plot visualizations, while schematic diagrams were prepared in BioRender and Adobe Illustrator.

## Data availability

The raw and processed FULL-seq datasets generated in this study have been deposited in the NCBI Gene Expression Omnibus (GEO) under accession number GSE353800. The dataset includes matched single-cell RNA and ATAC profiles from pooled human and mouse cell lines. The GEO records are currently private and will be made publicly available upon publication.

Publicly available RNA-sequencing datasets used for external validation were obtained from the NCBI Gene Expression Omnibus (GEO) under accession numbers GSE138248 (parental and liver-metastatic colorectal cancer patient-derived xenografts) and GSE221240 (colorectal cancer patient tissues, including primary tumors and liver metastases). RNA-sequencing data from colorectal cancer cell lines in the Cancer Cell Line Encyclopedia (CCLE) were obtained from the NCBI Sequence Read Archive (SRA) under BioProject accession PRJNA523380 (SRA study SRP186687). The 38 CCLE samples included in the analysis, together with their cell-line identifiers, SRA run accessions and analysis-group assignments, are listed in Supplementary Table 3.

Reference genomes and gene annotations (mm10 and hg38) were obtained from the UCSC Genome Browser. All other relevant data supporting the findings of this study are available from the corresponding author upon reasonable request.

## Acknowledgements

We thank colleagues at the Cutaneous Biology Research Center for providing resources and support. This work was supported by NIH New Innovator Award (DP2TR004354, JS), NIH Common Fund (UH3CA275687, UG3CA275687, JS), NIH (P01AI129859, JS), Massachusetts Life Science Center (JS), Burroughs Wellcome Fund (JS), Additional Ventures (JS), and Massachusetts General Hospital (JS).

## Author contributions

Under the supervision of J.S., M.W. conceived and developed the FULL-seq method, designed the study, and performed all experiments. M.W. devised the research strategy and led the overall project. M.W., J.Q., J.S., F.M., X.C., and A.Q. performed the computational analyses. M.W., J.S., and L.Z. wrote the manuscript with input from all authors. M.W., J.S., L.Z., F.M., X.C., and A.Q. contributed to data interpretation and discussion. J.S. and L.Z. supervised the study and secured funding. All authors read and approved the final manuscript.

