## Extended Figures for "FULL-seq reveals isoform-specific regulatory architecture in single cells"

### Extended Data Fig. 1

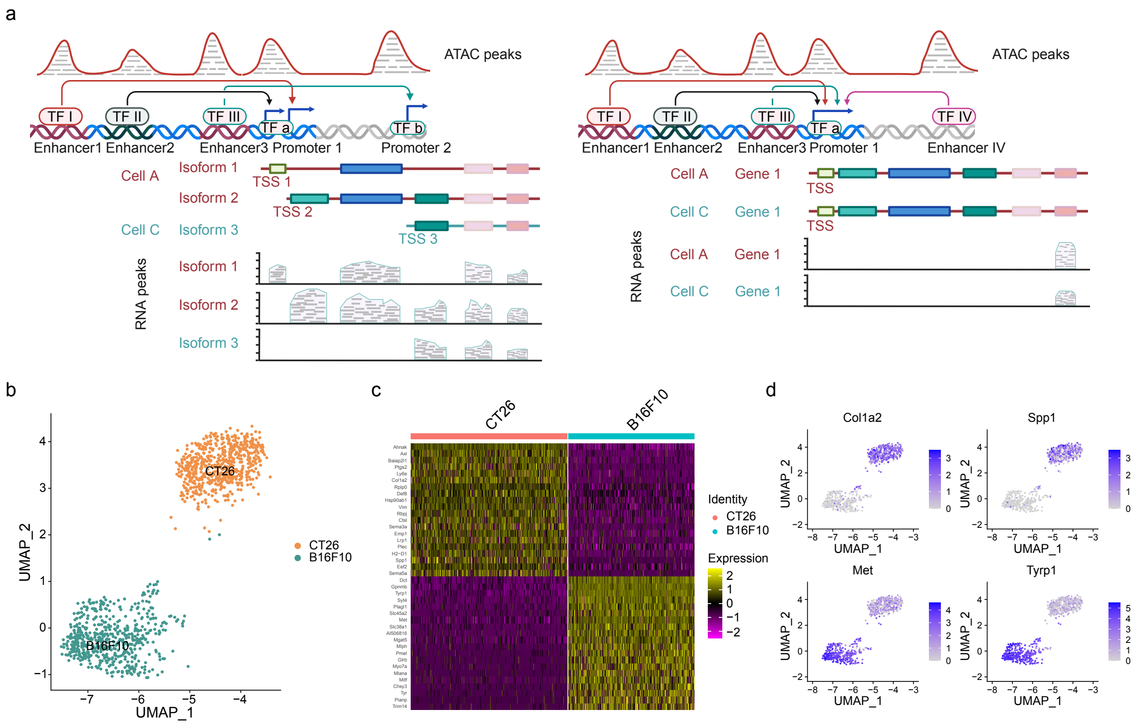

**Extended Data Fig. 1: Conceptual framework of isoform-resolved regulatory mapping and benchmarking of FULL-seq gene expression in a CT26 and B16F10 mixture.**

**a,** Comparison of regulatory mapping between FULL-seq (left) and conventional Multiome-seq (right). FULL-seq links ATAC peaks directly to isoform-specific TSSs, resolving enhancer–promoter–isoform relationships at single-cell resolution, whereas conventional methods lack isoform resolution, aggregating RNA signals to gene bodies and obscuring regulatory complexity.

**b,** UMAP visualization of FULL-seq gene expression profiles from a CT26 and B16F10 cell mixture. Cells are colored according to clusters identified from RNA expression profiles.

**c,** Heatmap showing the top differentially expressed genes distinguishing CT26 and B16F10 cells.

**d,** Feature plots of representative marker genes overlaid on the UMAP projection, demonstrating cluster-specific expression.

### Extended Data Fig. 2

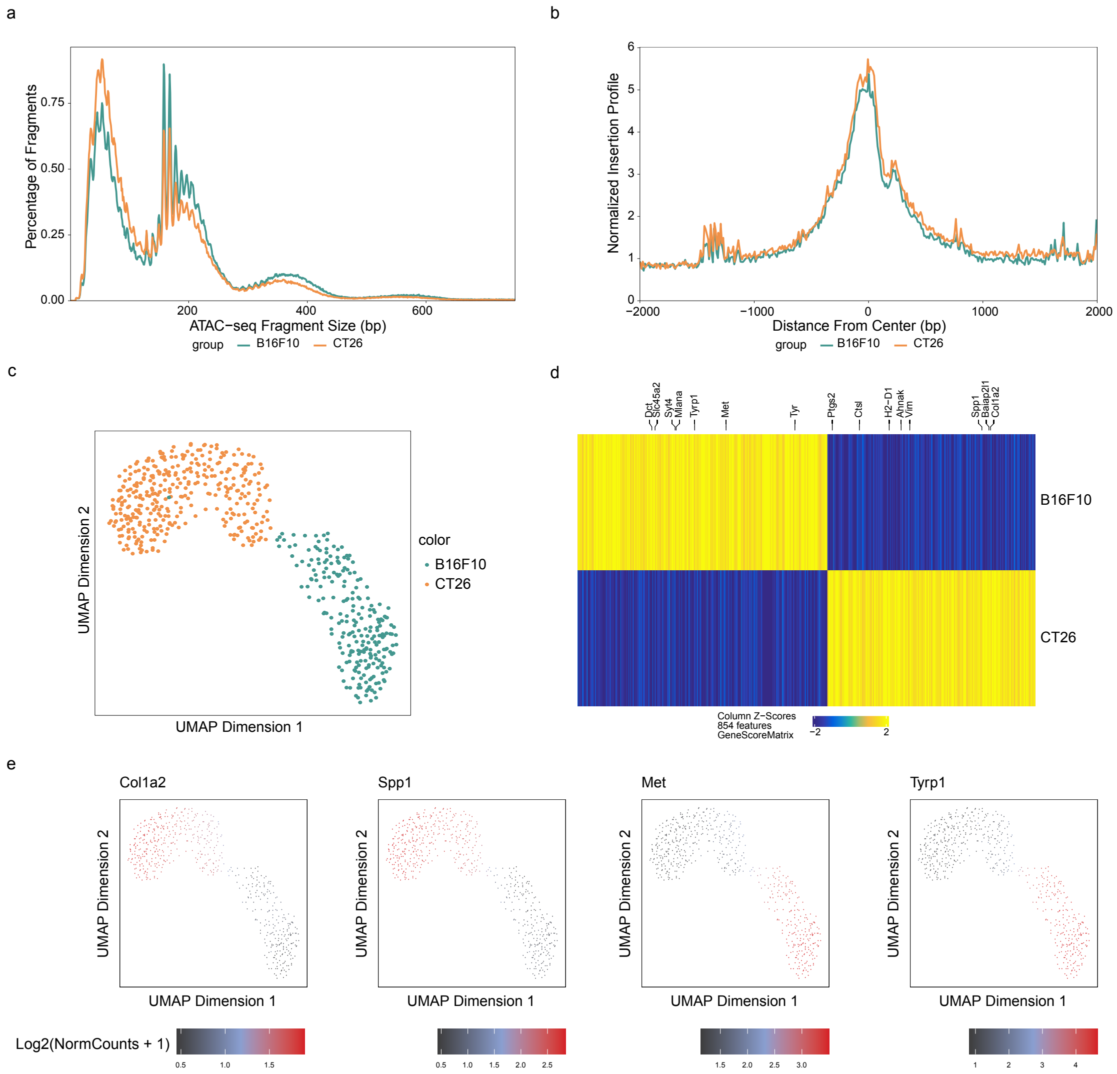

**Extended Data Fig. 2: Quality assessment and clustering of FULL-seq ATAC data from CT26 and B16F10 cells.**

**a,** Insert size distribution of ATAC-seq fragments, showing expected nucleosomal periodicity.

**b,** Aggregated ATAC-seq read enrichment around transcription start sites (TSSs).

**c,** UMAP visualization of FULL-seq ATAC profiles from a CT26 and B16F10 cell mixture. Cellular identities are colored based on independent clustering results with ATAC data.

**d,** Heatmap of differentially accessible genomic regions across clusters.

**e,** Feature plots showing gene activity scores (derived from ATAC data) for cell-type marker genes: Col1a2 and Spp1 for CT26, and Met and Tyrp1 for B16F10.

#### Extended Data Fig. 3

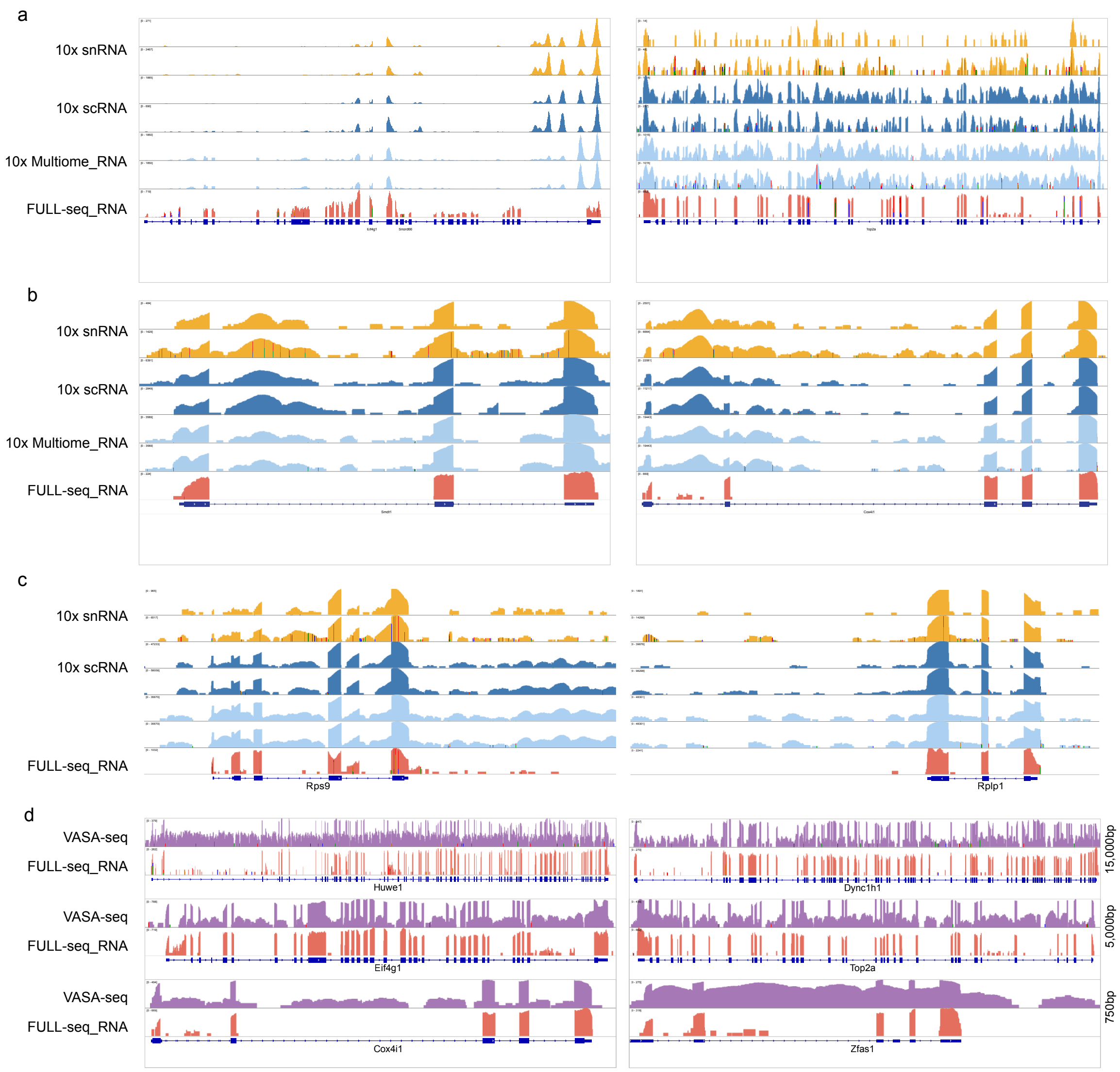

**Extended Data Fig. 3: Comparison of RNA coverage across platforms at representative loci.**

**a,** IGV genome browser tracks showing aggregated RNA coverage from single cells profiled by 10x snRNA-seq, 10x scRNA-seq, 10x Multiome-seq, and FULL-seq at representative loci spanning 5,000 bp (left) and 10,000 bp (right) regions.

**b,** IGV genome browser tracks showing aggregated RNA coverage from single cells profiled by 10x snRNA-seq, 10x scRNA-seq, 10x Multiome-seq, and FULL-seq at representative loci spanning 500 bp (left) and 750 bp (right) regions.

**c,** IGV genome browser tracks displaying aggregated RNA coverage from 10x snRNA-seq, 10x scRNA-seq, 10x Multiome-seq, and FULL-seq. Conventional platforms show increased intergenic signal compared to FULL-seq.

**d,** IGV genome browser tracks comparing aggregated RNA coverage from VASA-seq and FULL-seq, highlighting elevated intronic signal in VASA-seq relative to FULL-seq.

### Extended Data Fig. 4

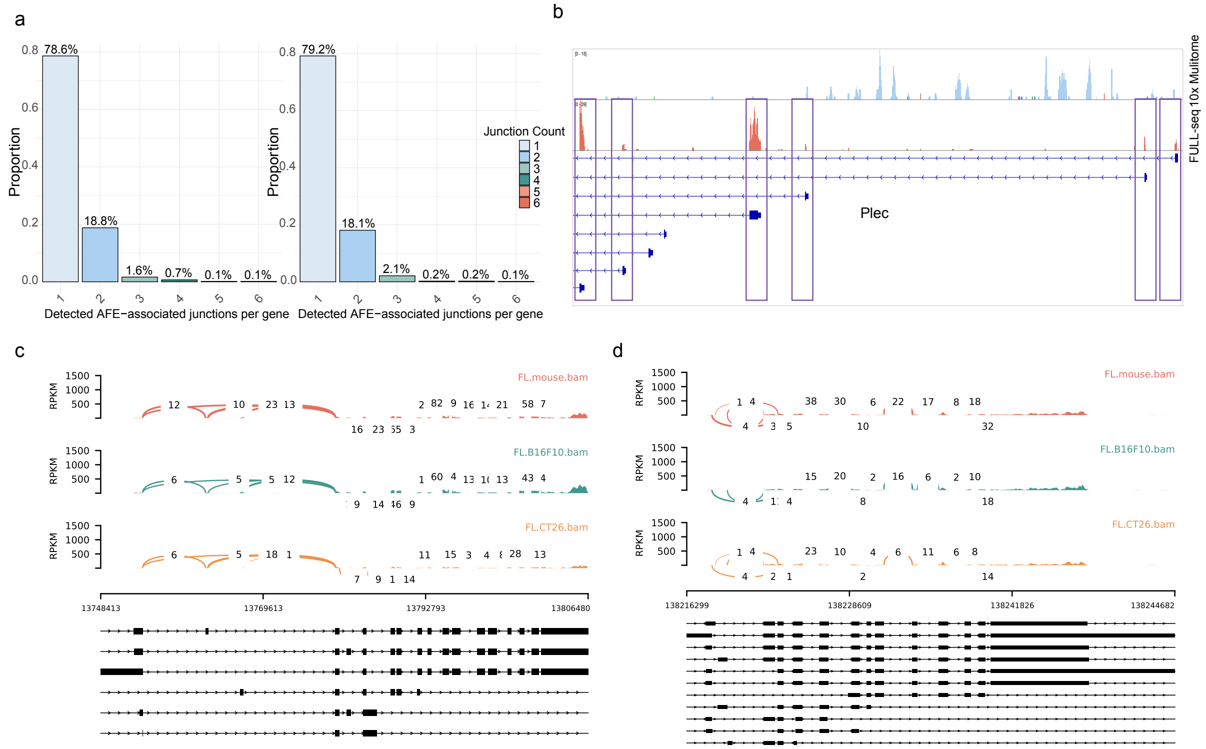

**Extended Data Fig. 4: Distribution and examples of alternative first exon (AFE) usage in CT26 and B16F10 cells.**

**a,** Distributions of the number of distinct detected AFE-associated junctions per gene in B16F10 (left) and CT26 (right) cells, analyzed separately using FULL-seq RNA data.

**b,** IGV genome browser tracks illustrating six distinct annotated AFEs within a representative gene, with RNA coverage and splice junction evidence supporting alternative first-exon expression.

**c–d,** Sashimi plots illustrating genes expressing three (c) and four (d) AFEs across combined, B16F10, and CT26 cells using FULL-seq RNA data. Read density is represented by bar heights, and splice junction usage by arcs, with read counts labeled.

### Extended Data Fig. 5

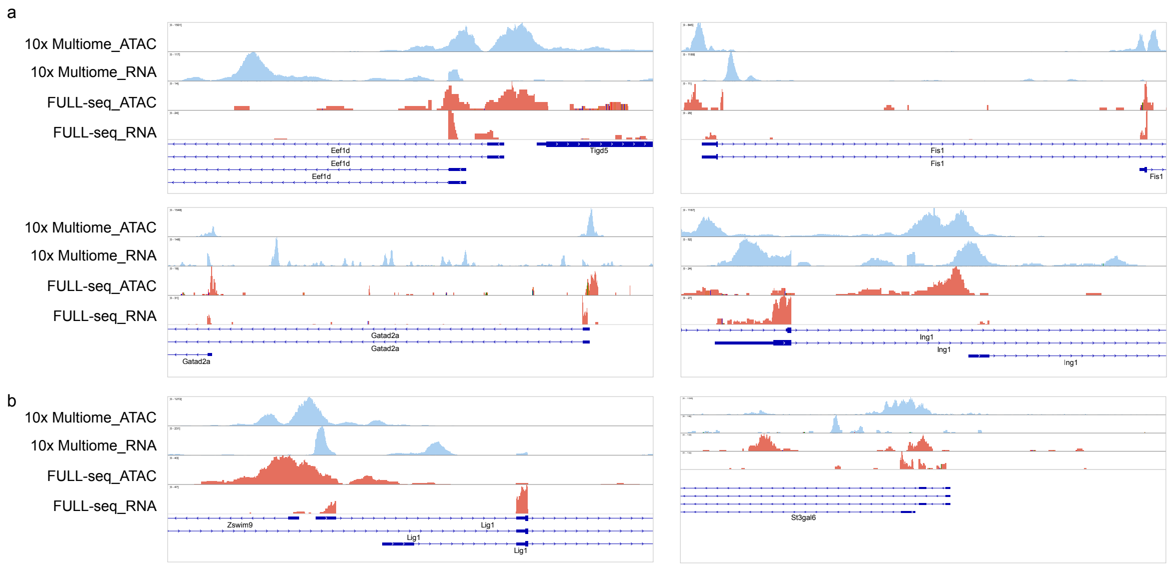

**Extended Data Fig. 5: Divergent relationships between AFE usage and chromatin accessibility across loci.**

a, IGV genome browser tracks showing chromatin accessibility and AFE expression across representative genes. The top panels highlight loci where ATAC peak intensities are comparable, yet AFE expression levels differ substantially. The bottom panels illustrate regions where chromatin accessibility and AFE expression vary independently, highlighting heterogeneous relationships between promoter accessibility and AFE expression.

b, IGV tracks of loci containing two adjacent genes. The left panels depict cases where neighboring genes share an ATAC peak but differ in expression, with one gene expressed at high levels and the other at low levels. The right panels show instances where a shared ATAC peak exists, yet only one gene is transcriptionally active.

### Extended Data Fig. 6

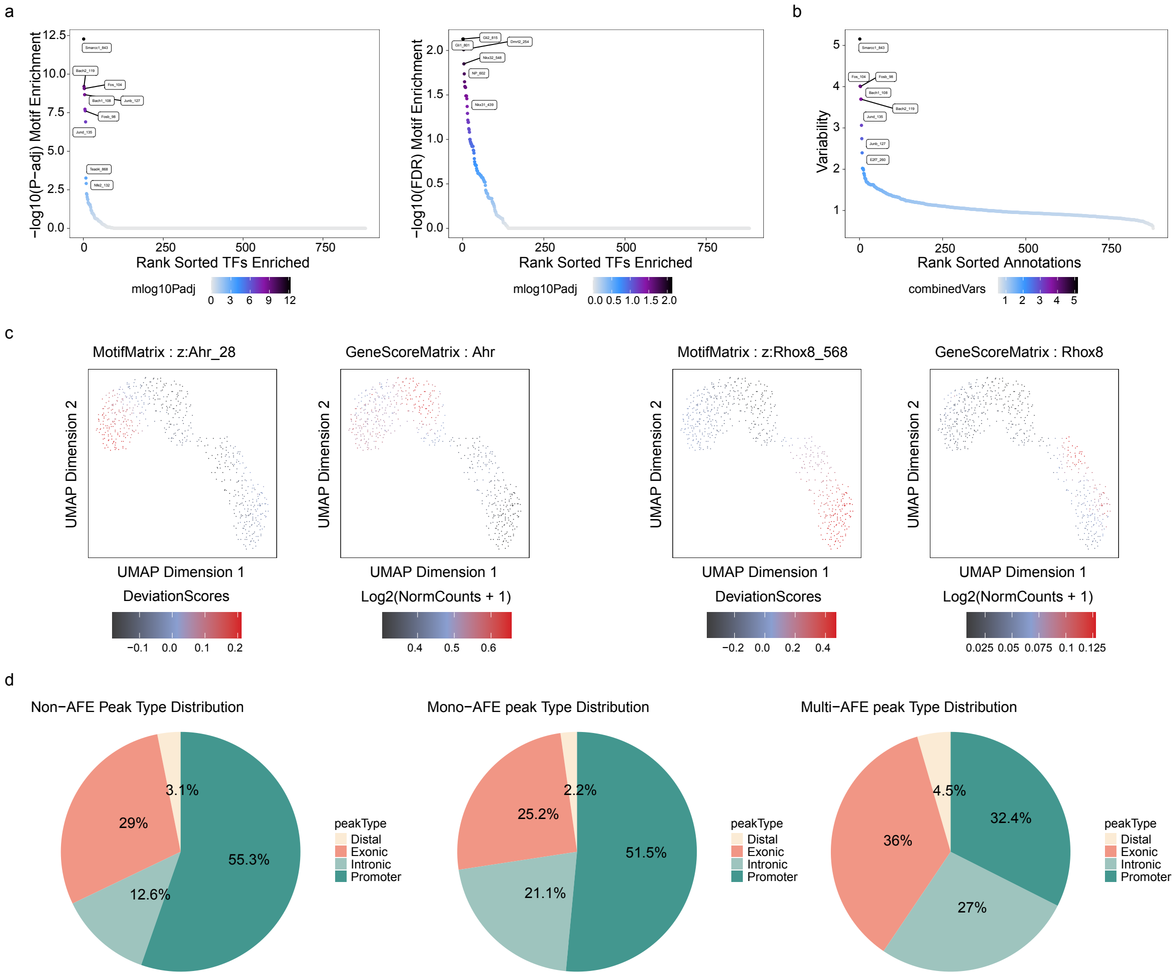

**Extended Data Fig. 6: Chromatin accessibility and transcription factor motifs reveal distinct regulatory features of AFE usage in CT26 and B16F10 cells.**

a, Rank-sorted motif enrichment analysis of transcription factors (TFs) in CT26 (left) and B16F10 (right) cells, with color scale indicating adjusted p-values (mlog10Padj). Darker colors represent stronger motif enrichment.

b, ChromVAR analysis of TF motif deviations, ranked by deviation scores, highlighting highly variable motifs across cells.

c, UMAP visualizations showing motif deviation scores (MotifMatrix) and gene activity (GeneScoreMatrix) for representative TFs Ahr and Rhox8, demonstrating correlated motif activity and gene expression patterns.

d, Pie charts showing the distribution of conventional genomic annotations of ATAC peaks overlapping regions associated with non-AFE, mono-AFE, and multi-AFE genes. Peaks were classified as promoter, exonic, intronic, or distal using ArchR's gene-based annotation framework.

### Extended Data Fig. 7

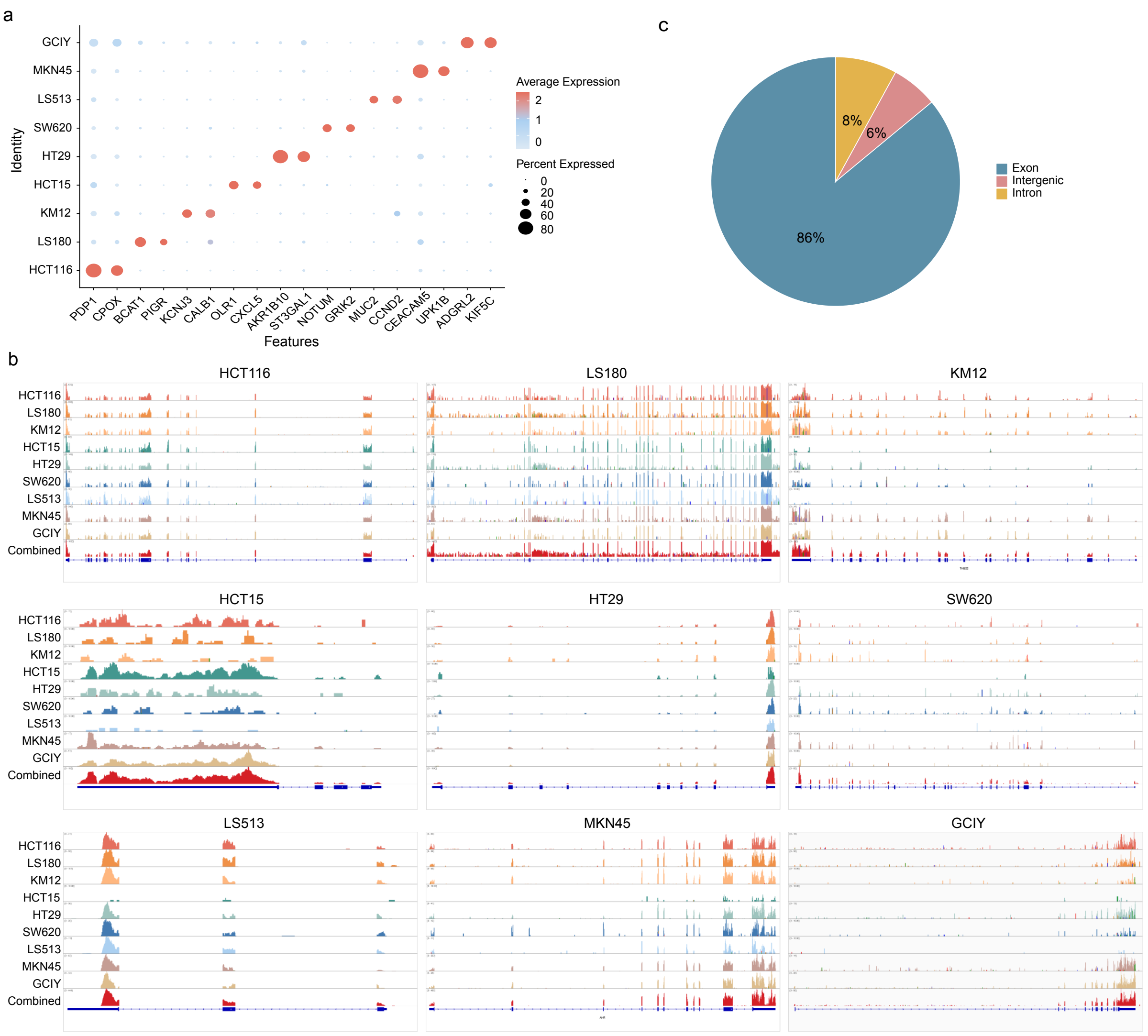

**Extended Data Fig. 7: FULL-seq captures gene expression patterns and isoform-level transcript coverage across nine human cancer cell lines.**

a, Dot plot showing average expression (color scale) and the percentage of cells expressing each marker gene (dot size) across nine cancer cell lines based on FULL-seq RNA profiles, confirming accurate cell-type identification.

b, IGV genome browser tracks displaying aggregated full-length mRNA coverage for selected marker genes across each cancer cell line. Even for genes with relatively low read counts, transcripts map cleanly to annotated exons, illustrating FULL-seq's ability to achieve isoform-resolved coverage even under sparse transcriptional input.

c, Fraction of reads mapped to exonic, intronic, and intergenic regions across the nine cell lines, showing the high exonic capture efficiency of FULL-seq.

### Extended Data Fig. 8

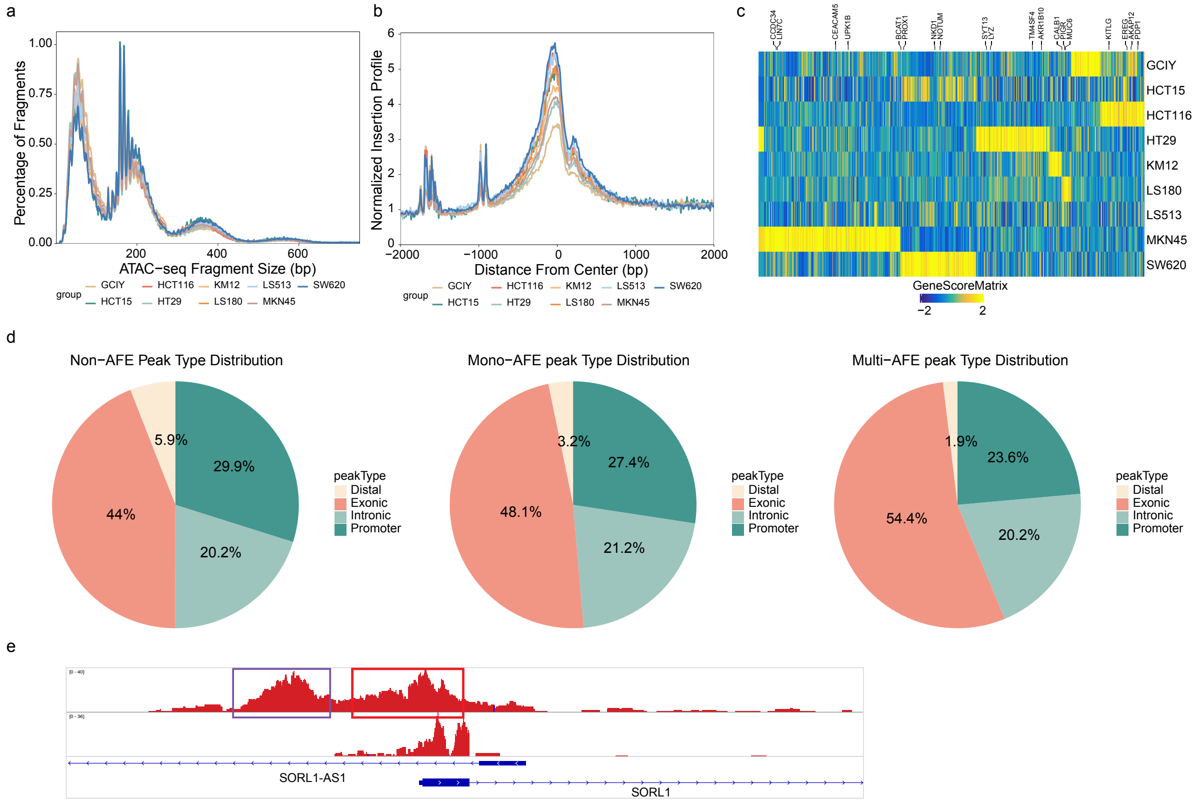

**Extended Data Fig. 8: FULL-seq reveals chromatin accessibility and AFE isoform-resolved transcriptomic features across cancer cell lines.**

a, Insert size distribution of ATAC-seq fragments across cell lines, showing expected nucleosomal periodicity.

b, Aggregate enrichment of ATAC-seq reads around transcription start sites (TSSs), confirming high data quality across samples.

c, Heatmap showing differential gene activity across nine cancer cell lines, based on chromatin accessibility-derived gene activity scores calculated using ArchR.

d**,** Pie charts showing the distribution of conventional genomic annotations of ATAC peaks associated with non-AFE, mono-AFE, and multi-AFE genes in human cells. Peaks were annotated as promoter, exonic, intronic, or distal using ArchR's gene-based annotation framework.

e, IGV browser view of representative loci where the AFE TSS (purple box) overlaps directly with a proximal ATAC peak. Many of these TSSs also fall within ±2,000 bp of additional distal peaks (red box), which are counted as “promoter-associated” in broad-window analyses.

### Extended Data Fig. 9

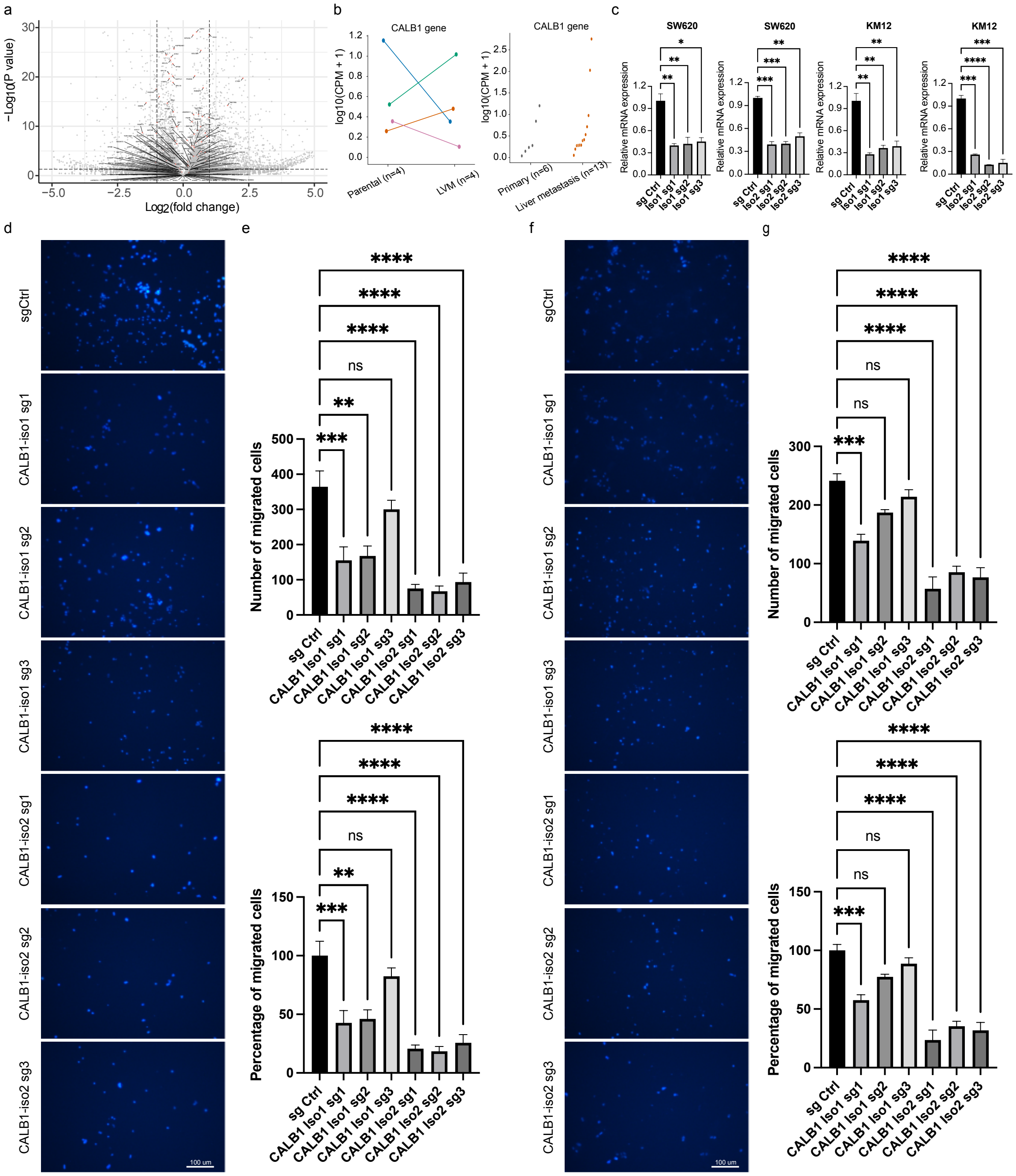

**Extended Data Fig. 9: Gene-level analysis and individual sgRNA validation of metastasis-associated AFE regulation.**

**a,** Volcano plot showing differential gene expression between primary and metastatic colorectal cancer cell lines detected by FULL-seq; labeled genes correspond to genes harboring top metastasis-associated AFE candidates.

**b,** Gene-level CALB1 expression in GSE138248 (left) and GSE221240 (right), corresponding to the AFE-level analyses in Fig. 5c,d. Matched parental and liver-metastatic (LVM) derivatives in GSE138248 (n = 4 pairs) are connected by lines; primary tumors (n = 6) and liver metastases (n = 13) in GSE221240 are shown as individual samples. Values are log10(CPM + 1).

**c,** RT–qPCR validation of CRISPRi-mediated repression of the targeted *CALB1* AFE. Expression of the targeted *CALB1* AFE was measured to assess knockdown efficiency following CRISPRi.

**d,** Representative Transwell migration images of SW620 cells following CRISPRi-mediated repression of CALB1 Iso1 or Iso2 using three independent sgRNAs for each isoform. Cells expressing a control sgRNA (sgCtrl) were included as controls.

**e,** Quantification of the Transwell migration assays shown in **d**, presented as the number of migrated cells (top) and percentage relative to control (bottom).

**f,** Representative Transwell migration images of KM12 cells following CRISPRi-mediated repression of CALB1 Iso1 or Iso2 using three independent sgRNAs for each isoform.

**g,** Quantification of the Transwell migration assays shown in **f**, presented as the number of migrated cells (top) and percentage relative to control (bottom).

Data are presented as mean ± s.e.m. from 3 independent experiments. Statistical significance was determined by one-way ANOVA followed by Dunnett's multiple-comparisons test versus control. ***P** < 0.05, ****P** < 0.01, *****P** < 0.001, ******P** < 0.0001; ns, not significant.
