## Supplementary Figures for "FULL-seq reveals isoform-specific regulatory architecture in single cells"

### Supplementary Fig.1

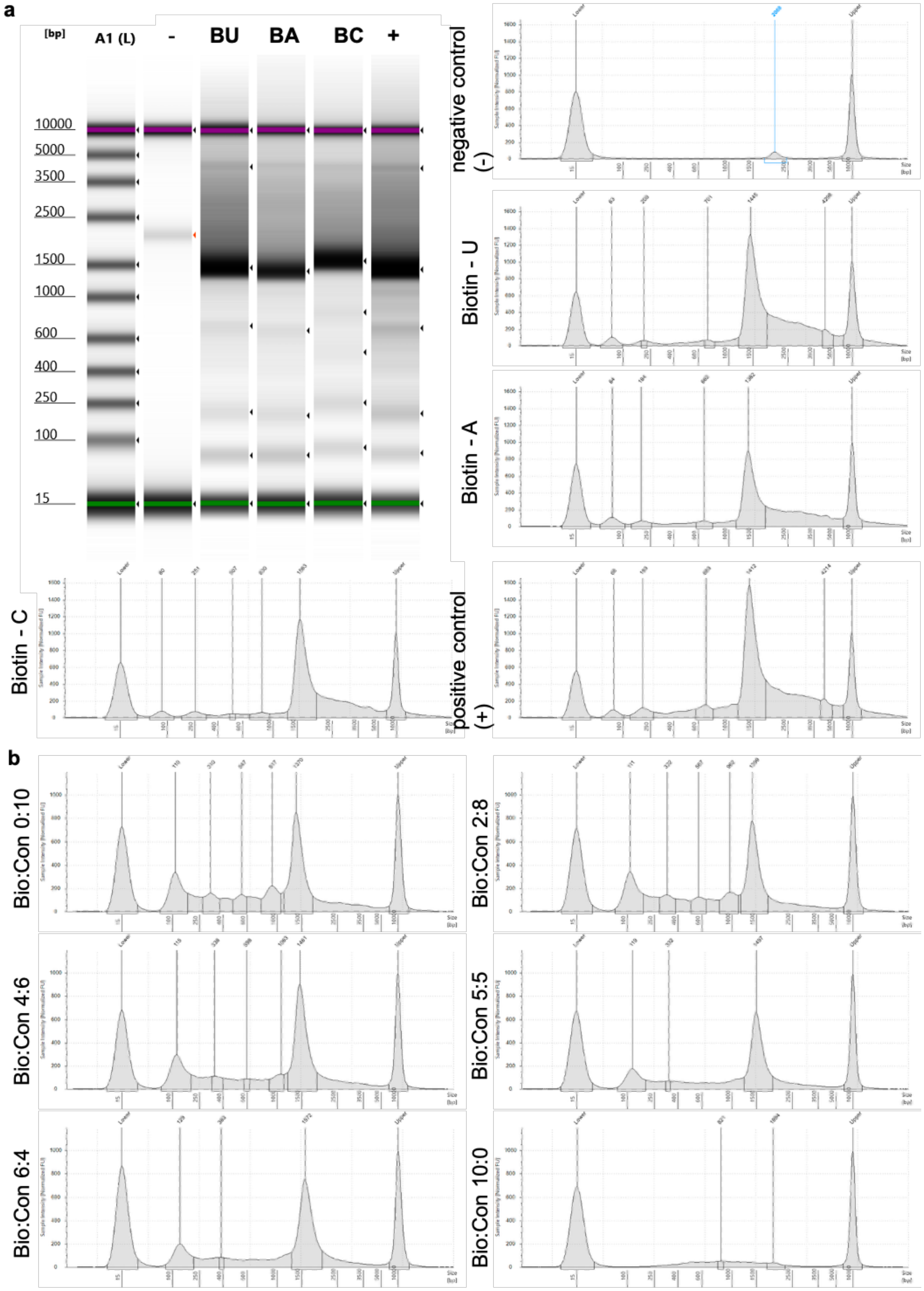

**Supplementary Fig.1: Evaluation of reverse transcription efficiency using biotinylated dNTP.**

**a,** Bioanalyzer electropherogram gel (left) and corresponding traces (right) showing cDNA size profiles from reverse transcription reactions performed with standard primers and supplemented with biotinylated dUTP (BU), biotinylated dATP (BA), or biotinylated dCTP (BC). Lane "–" denotes a no-primer negative control, and "+" is a positive control using standard oligo(dT)-primed reactions without biotinylated nucleotides. All three biotinylated dNTPs supported robust cDNA synthesis, yielding strong signals in the 1,000–3,000 bp range, while the negative control showed minimal background and the positive control displayed the expected full-length cDNA distribution.

**b,** Electropherogram traces of cDNA generated with Biotin-dCTP as the biotin source under different nucleotide mixing ratios. Standard dCTP and biotin-dCTP were combined at varying proportions to assess their effects on reverse transcription efficiency. Each panel represents a separate condition, with clear cDNA peaks (800–2,000 bp) indicating successful full-length cDNA synthesis.

### Supplementary Fig. 2

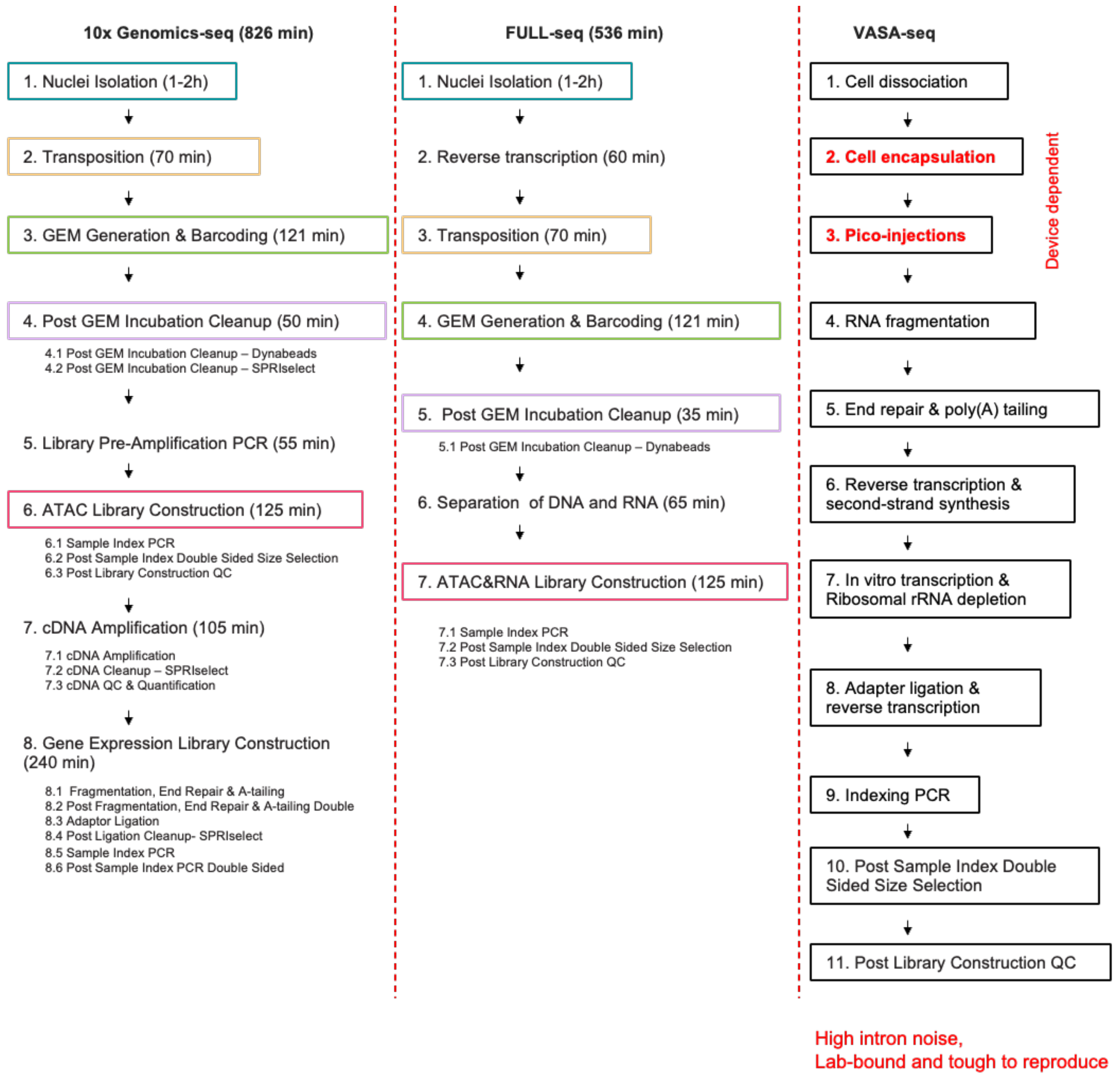

**Supplementary Fig. 2: Workflow comparison of FULL-seq, 10x Genomics-seq, and VASA-seq for single-cell multi-omics profiling.**

This schematic compares processing time, workflow complexity, and key steps across three protocols: 10x Genomics-seq (left, total ~826 min), FULL-seq (middle, ~536 min), and VASA-seq (right, time not specified). FULL-seq performs reverse transcription and tagmentation before droplet-based barcoding, followed by RNA–DNA separation and independent library construction. By contrast, 10x Genomics-seq requires separate library preparations for ATAC and gene expression, extending the overall runtime and necessitating additional PCR amplification. VASA-seq includes device-dependent steps such as cell encapsulation and pico-injection, making it less reproducible and contributing to elevated intronic signal noise, as observed in IGV comparisons.

### Supplementary Fig. 3

**a**

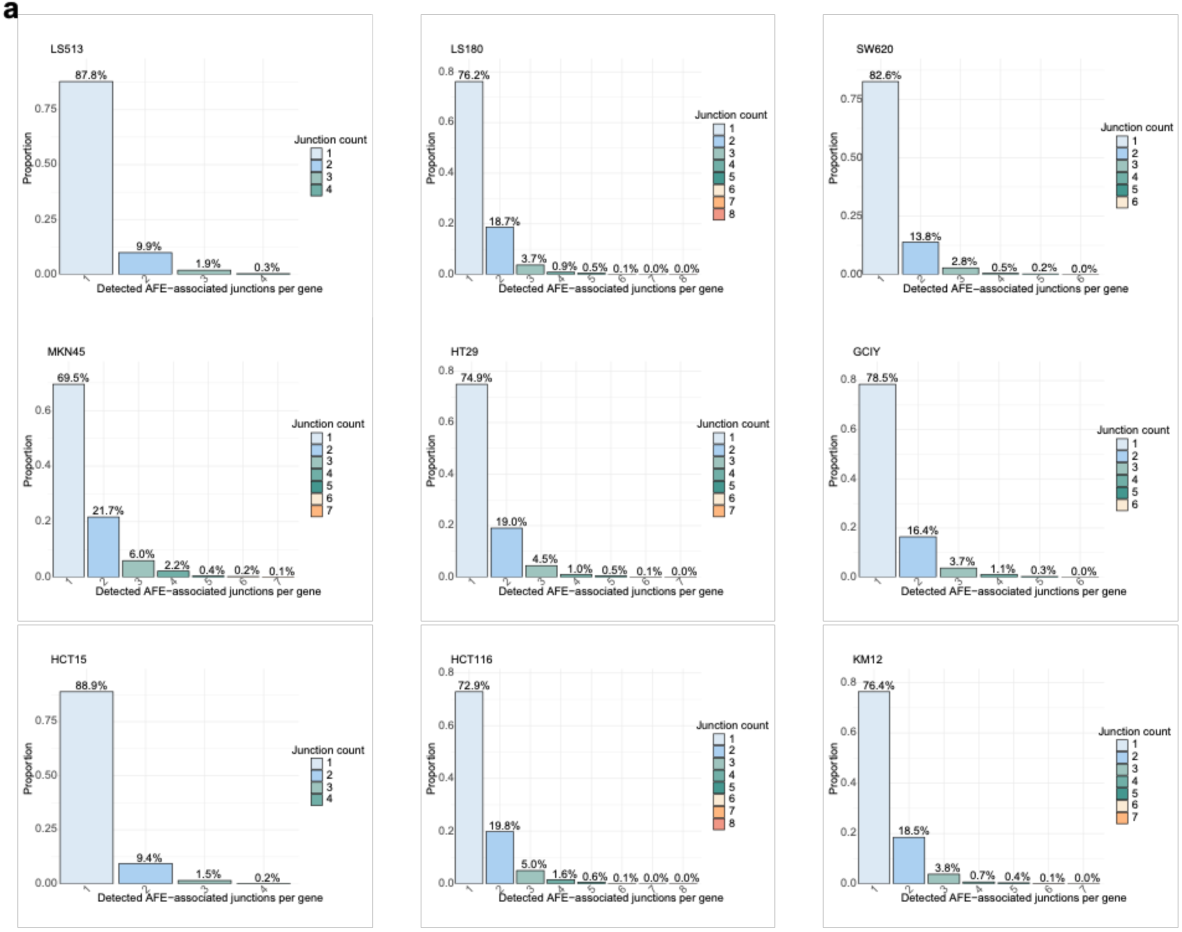


**b**

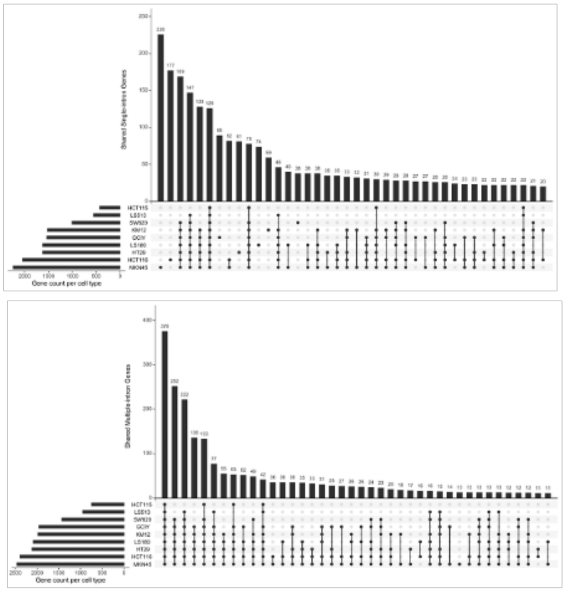

**c**

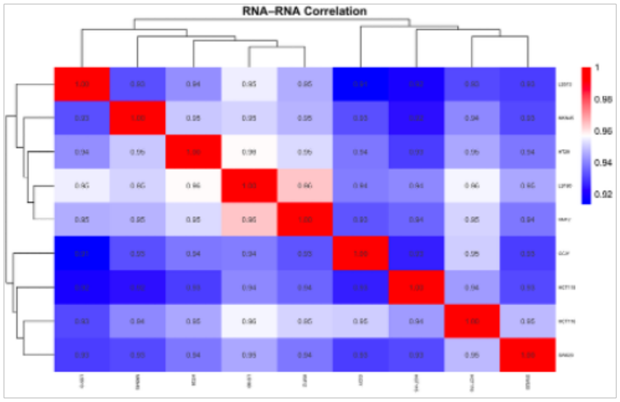

**Supplementary Fig. 3: Distribution and cross-line overlap of AFE usage across nine human cancer cell lines.**

**a,** Bar plots showing the distributions of distinct detected AFE-associated junctions per gene across nine human cancer cell lines. Each cell line was analyzed independently, and genes were counted once according to the number of distinct junctions detected. The distributions illustrate variation in AFE-associated junction multiplicity across cell lines.

**b,** UpSet plots showing the overlap of genes with a single or multiple detected AFE-associated junctions across the nine cell lines. Horizontal bars indicate the number of genes in each cell line, whereas vertical bars represent the number of genes shared among the indicated cell-line intersections.

**c,** Correlation heatmap of gene-level expression across nine human cancer cell lines, providing a reference for comparison with the AFE-level correlation analysis shown in Fig. 4e.
